# Pubertal maturation and chemotherapy-associated disruption of the pediatric ovary revealed by multimodal single-cell profiling

**DOI:** 10.64898/2026.06.03.729786

**Authors:** Loren Méar, Jasmin Hassan, Matthew William Myers, Hosein Toosi, Ilmatar Rooda, Nina Boskovic, Filippa Bertilsson, Feria Hikmet, Anastasios Damdimopoulos, Rutger Schutten, Borbala Katona, Marzie Abdolhamdi, Emmanouela Perisynaki, Katri Knuus, Karin Pettersson, Kiriaki Papaikonomou, Johan Malmros, Petra Byström, Mikael Sundin, Cecilia Langenskiöld, Hartmut Vogt, Géraldine Giraud, Andres Salumets, Marjut Otala, Timo Tuuri, Joakim Lundeberg, Andrea Jurisicova, Cecilia Lindskog, Kirsi Jahnukainen, Reza Mirzazadeh, Pauliina Damdimopoulou

**Author notes:** Shared first authorship (Loren Méar, Jasmin Hassan, Matthew William Myers). Shared senior authorship (Kirsi Jahnukainen, Reza Mirzazadeh, Pauliina Damdimopoulou). Corresponding author: Loren Méar, Kirsi Jahnukainen, Reza Mirzazadeh, Pauliina Damdimopoulou.

## Abstract

Ovarian tissue cryopreservation enables fertility preservation in females undergoing gonadotoxic therapies, restoring fertility in adults. Although offered even before puberty, the childhood ovary and its vulnerability to therapy remain poorly characterized. Here, ovarian tissue from 16 patients undergoing fertility preservation (aged 1-16 years) and 11 adult controls (aged 22-32 years) was analyzed using single-cell RNA sequencing, spatial transcriptomics, and multiplex immunostaining. In chemotherapy-naïve samples, 13 somatic cell populations underwent extracellular matrix remodeling, vascular, neural, and stromal maturation during puberty, whereas changes in germline related to chromatin remodeling. Spatial transcriptomics resolved 23 clusters across, revealing distinct tissue organization and follicular niche composition between children and adults. Chemotherapy exposure depleted perifollicular and vascular cells, suppressed intercellular signaling, and dysregulated over half of puberty-associated genes, converging on stress responses and extracellular matrix remodeling, with SEPTIN7 as a potential biomarker. These findings uncover critical developmental vulnerabilities of the pediatric ovary relevant to fertility preservation.

## INTRODUCTION

Fertility preservation aims to cryostore gametes in patients undergoing gonadotoxic treatments and at high risk of infertility. In postpubertal females, the standard method is vitrification of mature oocytes following hormonal stimulation (*1*). When treatment cannot be delayed, ovarian tissue cryopreservation (OTC) can instead be performed (*1*). Introduced in the 1990s, OTC followed by autotransplantation has resulted in over 200 live births worldwide (*2*), with reported live birth rates of 21–33% in adults, approaching those achieved with *in vitro* fertilization in this setting (*3*).

OTC preserves a portion of the ovarian reserve, comprising dormant follicles located in the ovarian cortex. The reserve is established during fetal development, when primary oocytes arrest in prophase I to form primordial follicles, and declines with age through atresia and growth activation (*4, 5*). Although full maturation and ovulation occur only after puberty, follicles undergo gonadotropin-independent growth to the secondary stage already during childhood (*4, 6*). Accordingly, both pediatric and adult ovarian cortex contain primordial, primary, and secondary follicles, supporting the use of OTC in prepubertal girls with the expectation that retransplanted tissue may later restore fertility (*7, 8*).

Puberty in girls spans approximately 1.5–6 years and typically begins between 9 and 14 years of age (*9*). Ovarian volume increases roughly tenfold from birth to adulthood (*10*), accompanied by altered cortical mechanics, with a transition from a rigid extracellular matrix (ECM) in childhood to increased viscoelasticity in adults (*11*). Concurrently, the ovarian reserve and the proportion of morphologically abnormal follicles decline (*5, 12*). Compared to adults, follicles from prepubertal ovaries show reduced *in vitro* growth capacity (*12, 13*) and distinct transcriptomic profiles (*14*). Evidence on chemotherapy-induced damage in children remains limited to histological observations of increased follicular atresia and *in vitro* studies showing reduced growth and steroidogenesis (*15, 16*). Despite recent single-cell ovarian atlases, pediatric tissue remains underrepresented (*17*), and systematic comparisons across age and treatment exposure are lacking.

To date, no live births have been reported following transplantation of ovarian tissue cryopreserved in early childhood. Only three pregnancies using pediatric tissue have been described, all in patients of pubertal age (9, 14, and 14 years) at tissue harvest (*18–20*). The use of OTC in young children without evidence of future fertility restoration therefore raises important clinical and ethical considerations.

Here, ovarian tissue collected through a clinical pediatric fertility preservation study was analyzed together with adult samples to define ovarian maturation and the effects of chemotherapy exposure. Analysis of 27 samples revealed extensive stromal remodeling across puberty, accompanied by vascular and neural maturation and a shift in germline transcriptional programs. Chemotherapy induced broader transcriptional alterations than puberty, including ECM disruption and reduced intercellular communication. These findings challenge the direct extrapolation of adult OTC protocols to prepubertal patients.

## RESULTS

### Study design and cohort characterization

The success of fertility preservation depends on the quality of the cryopreserved ovarian cortex, clinically assessed by follicle morphology (**Fig. 1a**) (*21, 22*). Haematoxylin–eosin (H&E) staining shows dense stroma with scattered primordial follicles in adults, whereas pediatric samples display looser stroma with higher follicle density (**Fig. 1a**). However, histology provides limited insight into tissue maturity and treatment effects. Higher-resolution profiling may enable biomarker discovery to improve quality assessment and patient selection for OTC.

**Figure 1.**
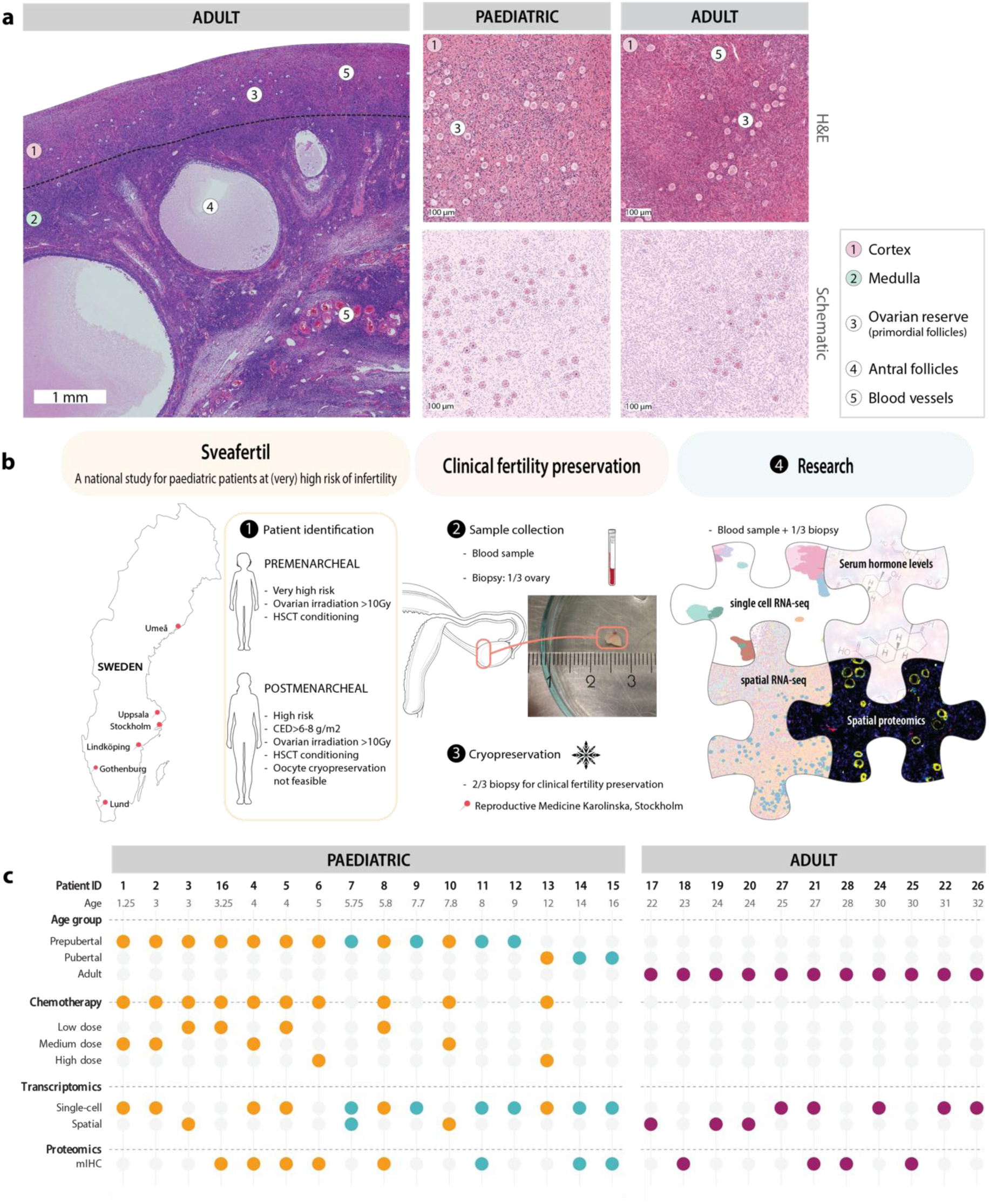
Study design and cohort characterization. ***a, Representative H&E–stained ovarian tissue sections.*** Left, low-magnification overview of an adult ovary showing the (1) cortex (approximately 1 mm in depth) and the (2) medulla, separated by a dashed line. The cortex contains the (3) ovarian reserve, composed of primordial follicles, whereas (4) antral follicles are located in the medulla rich in (5) blood vessels. Right, ovarian cortex from pediatric and adult ovaries. Pediatric samples are characterized by a looser stromal architecture and a higher density of (3) primordial follicles compared with adults. Schematic representations illustrate age-dependent differences in follicle and stroma density. ***b***, ***Study outline.*** Within the national pediatric fertility preservation program (Sveafertil), (1) eligible patients aged 1–17 years were identified across Sweden according to national inclusion criteria. (2) Peripheral blood and an ovarian biopsy (∼ one-third of one ovary) were collected. (3) Two-thirds of the biopsy was cryopreserved for clinical use at Reproductive Medicine in Karolinska University Hospital (Stockholm), and (4) one-third was allocated for research. Downstream analyses included serum hormone measurements, single-cell and spatial transcriptomics, and spatial proteomics. ***c, Cohort overview.*** Ovarian tissue from 16 pediatric patients (1–16 years) and 11 reproductive-age adults (22–32 years; dark fuchsia) was included for molecular profiling. Transcriptomic analyses comprised single-cell RNA sequencing and high-definition spatial transcriptomics, and proteomic profiling was performed using multiplex immunohistochemistry (mIHC). Pubertal status (prepubertal or pubertal) was assigned based on age, clinical assessment and circulating hormone levels. In pediatric patients, prior chemotherapy exposure is indicated (orange), and absence of exposure is shown in teal. Alkylating chemotherapy exposure was stratified by cumulative CED: low (<4000), medium (≥4000 and < 6000), and high (≥6000).

A national fertility preservation study was established for OTC in patients aged 1–17 years in accordance with Nordic and Swedish recommendations (sveafertil.ki.se) (**Fig. 1b**) (*23*). Eligibility criteria included hematopoietic cell transplantation or ovarian irradiation >10 Gy irrespective of pubertal status; in postpubertal patients, high-risk classification additionally included cumulative alkylating treatment [cyclophosphamide equivalent dose (CED) >6–8 g/m²]. Approximately one-third of one ovary was excised, of which two-thirds were cryopreserved for clinical use and one-third allocated for research (**Fig. 1b**).

In total, ovarian tissue from 16 pediatric patients aged 1–16 years was included. Diagnoses ranged from non-malignant hematological disorders (*n* =3) to leukemia (n=4) and solid tumors (*n* = 9). Ten patients had received chemotherapy before OTC, including nine exposed to alkylating agents (CED 1–8 g/m2) (**Fig. 1c, Supplementary Table 1**). Serum hormone analyses and clinical assessment indicated that three patients were pubertal at sampling (12–16 years of age) (**Supplementary Table 1**). Ovarian cortex from 11 adults aged 22–32 years served as a reference. All samples were collected between 2020 and 2024 and directly processed for single-cell RNA sequencing, spatial transcriptomics and multiplex immunostaining (**Fig. 1c**).

### Cellular landscape of chemotherapy-naïve ovarian cortex across development

Single-cell ovary transcriptomic datasets spanning childhood to reproductive age are lacking. To establish a reference, chemotherapy-naïve pediatric and adult samples were integrated for cell type identification and annotation. Integration of six pediatric (6–16 years) and five adult (25–32 years) samples yielded 26 clusters (**Supplementary Fig. 1a**). After removal of clusters representing noise, 21 remained for annotation (**Fig. 2a**).

**Figure 2.**
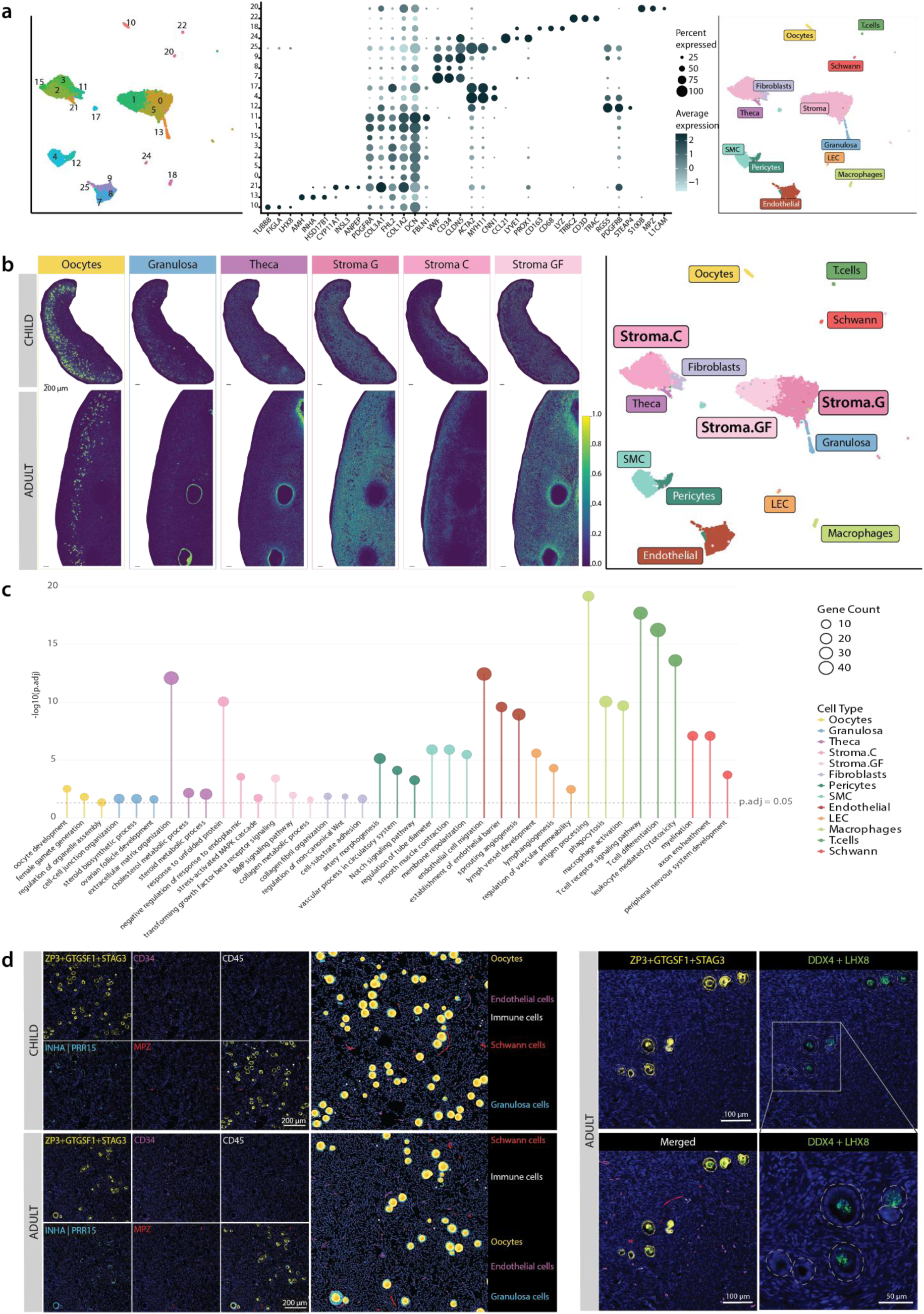
Cellular landscape of the chemotherapy-naïve human ovarian cortex across development. **a, Integrated single-cell RNA-sequencing analysis of six pediatric and five adult ovarian cortex samples.** Left, UMAP representation of the 21 clusters following integration and removal of noise. Middle, dot plot showing expression of three representative marker genes per cell type for annotation of major ovarian cell populations. Dot size indicates gene expression frequency, and color indicates average expression level. Right, UMAP colored by annotated cell types. **b, Spatial deconvolution analysis and refined cell type annotation**. Left, spatial transcriptomics-based deconvolution heatmaps illustrating the distribution of single cell clusters related to follicles (oocytes, granulosa cells, and theca precursor cells) and stroma (Stroma-G, Stroma-C, Stroma-GF) across child (top) and adult (bottom) tissue sections. Right, updated UMAP displaying the fourteen annotated cell types, including stromal subpopulations. **c, GO biological process enrichment analysis across annotated cell populations.** Lollipop plot showing selected enriched GO terms per cluster. Dot size represents gene count, colors indicate cell type, and lollipop height corresponds to –log10(adjusted P value). Dashed line indicates the significance threshold (adjusted P = 0.05). **d, mIHC validation of key cortical cell compartments**. Left, representative images of the multiplex panel targeting five cell populations (oocytes, granulosa cells, endothelial cells, immune cells, and Schwann cells) in both adult and child samples, alongside a schematic derived from the corresponding images. Right, example of panel flexibility showing incorporation of additional markers (DDX4 and LHX8) to distinguish two follicle subtypes within the same tissue context in an adult sample. Follicles are circled in grey, Type 1 follicles are positive for DDX4 and LHX8 (green staining).

Cell identities were assigned using differential gene expression with gene ontology (GO) enrichment, comparison with published datasets and Human Protein Atlas data, automated annotation (SingleR and large language models), and spatial transcriptomic deconvolution to confirm anatomical localization. Although automated approaches were limited by the scarcity of ovary-specific references, combined manual and spatially informed annotation resolved fourteen major cortical cell populations.

Previously described follicular, vascular, and immune cell types were readily identified based on canonical markers and localization (**Fig. 2a,b, Supplementary Table 2**) (*17*). Oocytes (*TUBB8, FIGLA, LHX8*) were distributed throughout the cortex in both pediatric and adult samples and were surrounded by granulosa cells (*AMH, INHA, HSD17B1*). Theca precursor cells (*CYP11A1, INSL3, ANPEP*) associated with growing follicles were more prominent in adult tissue (**Fig. 2b**). Vascular and lymphatic clusters comprised endothelial (*VWF, CD34, CLDN5*), smooth muscle (*ACTA2, MYH11, CNN1*), pericyte (*RGS5, PDGFRB*), and lymphatic endothelial (*CCL21, LYVE1, PROX1*) populations. A fibroblast cluster was defined by *PDGFRA* and *FBLN1* expression and enrichment of ECM genes (*DCN, COL1A2*) (**Fig. 2a, Supplementary Table 2**).

Canonical markers further identified immune and neural-associated populations, including macrophages (*CD163, CD68, LYZ*), T cells (*TRBC2, CD3D, TRAC*), and Schwann cells (*S100B, MPZ, L1CAM*). Schwann cell identity was confirmed by co-localization of MPZ with S100B and neuronal markers (MAPT, PRPH) (**Supplementary Fig. 1b**).

Stromal cells, representing the largest population, segregated into three spatially distinct subtypes: broadly distributed stroma-general (stroma-G), cortex-enriched stroma-cortex (stroma-C), and growing follicle-associated stroma (stroma-GF) (**Fig 2b, Supplementary Fig. 1c**).

GO enrichment supported the annotations (**Fig. 2c, Supplementary Table 3**). Oocytes, granulosa, and theca cells were enriched for “oocyte development”, “ovarian follicle development” and “steroid metabolic process”, respectively. Endothelial cells, smooth muscle cells, and pericytes were enriched for “endothelial cell migration”, “smooth muscle contraction” and “artery morphogenesis”, whereas immune populations showed enrichment for “TCR signaling” (T cells), “antigen presentation” (macrophages) and “lymphangiogenesis” (lymphatic endothelial cells) (**Fig 2c**). Schwann cells were enriched for “myelination”, and fibroblasts for “collagen fibril organization”. While stroma-G showed no significant enrichment, stroma-C was enriched for “response to unfolded protein” and stroma-GF for “TGFβ receptor signalling”, indicating localised regulatory functions (**Fig 2c**).

To spatially validate key functional compartments, a multiplex immunohistochemistry (mIHC) panel was developed. The panel included ZP3 (cell surface), GTSE1 (cytoplasm), and STAG3 (nucleus) for oocytes; INHA or PRR15 for granulosa cells; MPZ for Schwann cells; CD34 for endothelial cells; and CD45 for immune cells. Staining confirmed vascular, neural, and immune components in both pediatric and adult cortex, frequently in proximity to follicles (**Fig. 2d**).

The panel was designed to accommodate additional markers for targeted analyses. As an example, antibodies against DDX4 and LHX8 were incorporated to distinguish recently described follicle subtypes with similar morphology but distinct molecular profiles(*14*), enabling discrimination of Type 1 (DDX4/LHX8-positive) and Type 2 (DDX4/LHX8-negative) follicles within cortical tissue (**Fig. 2d**). Together, the findings reveal cellular diversity not captured by conventional histology (**Fig 1a, Fig 2d**).

### Spatial architecture of child and adult ovaries

To further resolve the structural organization of the human ovary, three pediatric (3–8 years) and three adult (22–24 years) specimens were profiled using the Visium HD spatial transcriptomics platform (**Fig 1c**). Pediatric sections comprised mainly cortex, whereas adult sections included also medulla. Following segmentation and integration, ∼ 3 billion reads, ∼ 4.68 million spatial barcodes, and ∼ 1.96 million cells, including 1,163 oocytes, were retained for analysis.

Unsupervised clustering identified 23 transcriptionally distinct spatial clusters (Sp-Cl) (**Fig. 3a-b**). These represent spatially coherent tissue domains rather than cell types, each defined by characteristic gene expression reflecting local cellular composition. Regions were annotated based on differential expression, spatial localization, and GO enrichment, and grouped into seven compartments: follicle-associated, vascular, cortical, medullary, cortex–medulla interface, generally distributed, and immune-enriched (**Fig. 3a–e, Supplementary Fig. 2a–e, Supplementary Tables 4–5**).

**Figure 3.**
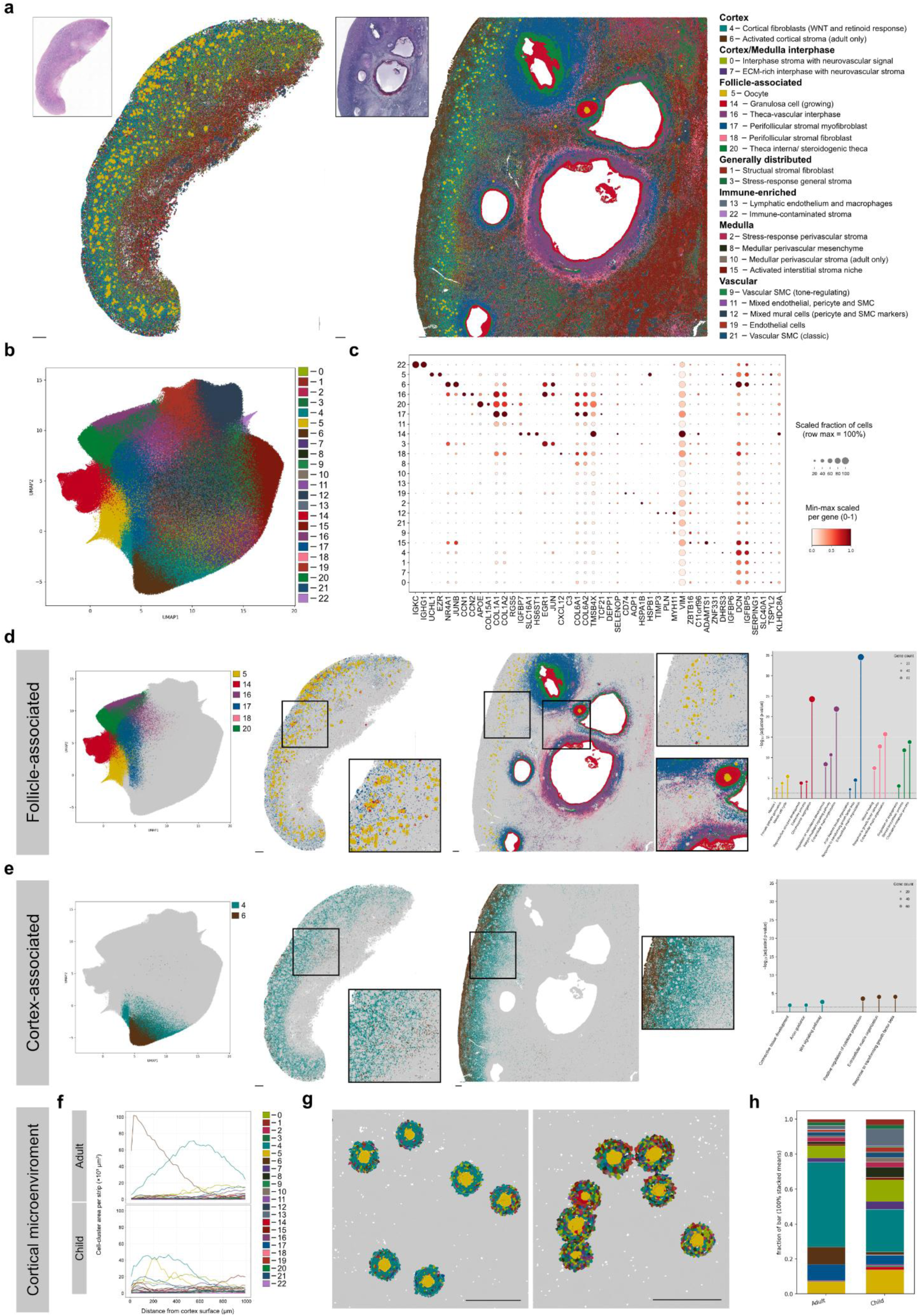
Spatial architecture of child and adult ovaries. **a, Spatial mapping of 23 transcriptionally distinct clusters onto representative tissue sections from a child (Patient 3, 3 years, left) and adult (Patient 17, 22 years, right), with corresponding H&E-stained images inset.** Clusters are grouped by spatial compartment in the legend: cortex, cortex/medulla interface, follicle-associated, generally distributed, immune-enriched, medullary, and vascular. **b, UMAP embedding of all 23 clusters identified jointly across all six samples.** **c, Dot plot of the top two marker genes per cluster.** Color intensity represents mean expression scaled between 0 and 1, and dot size represents the scaled fraction of cells expressing each marker (row maximum = 100%). **d, Follicle-associated compartment.** Left, UMAP highlighting follicle-associated clusters with remaining clusters greyed out. Middle, spatial localization on representative child and adult tissue sections, with magnified insets showing follicular structures. Right, lollipop plot showing GO enrichment; the y-axis shows −log₁₀(adjusted p-value) and dot size represents gene count. Dashed line indicates the significance threshold (adjusted P = 0.05). **e, Cortex-associated compartment.** Left, UMAP highlighting cortical stroma clusters with remaining clusters greyed out. Middle, spatial localization on representative child and adult tissue sections, with magnified insets. Right, lollipop plot showing GO enrichment, displayed as in (d). **f, Depth-resolved cortical composition.** Cell-cluster area per 20 µm strip plotted against distance from the cortical surface (outermost 1 mm) for adult (Patient 17, top) and child (Patient 3, bottom) representative sections, colored by cluster identity. **g, Perifollicular niche analysis.** Left, representative follicles from adult (Patient 17) and right, child (Patient 3) cortex illustrating the niche analysis approach: inner boundaries delineate oocyte borders and outer boundaries extend 30 µm beyond, defining the perifollicular ring region; cells within rings are colored by cluster identity. **h, Stacked bar plot showing mean fractional cluster composition within 30 µm perifollicular rings across the 137 follicles, stratified by maturity group (adult, child).** Scale bars: 200 µm throughout.

As expected, clear compartmentalization from surface to interior was observed(*24, 25*); the cortex was dominated by cortical fibroblasts (Sp-Cl4) with enriched WNT signaling, whereas medulla comprised perivascular and mesenchymal populations (Sp-Cl2, Sp-Cl8, Sp-Cl10) together with activated stroma (Sp-Cl15) enriched for stress response, ECM organization and signaling pathways (**Fig. 3a,e Supplementary Fig. 2e, Supplementary Table 5**). Two interphase regions (Sp-Cl0, Sp-Cl7) marked the cortex–medulla boundary (**Supplementary Fig. 2b**).

Follicular structures displayed layered organization, with oocyte (Sp-Cl5), granulosa (Sp-Cl14), and theca interna (Sp-Cl20) regions surrounded by perifollicular stromal domains enriched for ECM organization, vascular features, and contractility-associated transcripts (Sp-Cl16–18). Among these, perifollicular stromal myofibroblasts (Sp-Cl17) showed strong enrichment for ECM organization and expressed contractile markers *ACTA2* and *MYLK*, (**Fig. 3d, Supplementary Tables 4–5**). Although associated with antral follicles, Sp-Cl17 was also detected adjacent to cortical follicles in both age groups (**Fig. 3d**). As cortical stiffness regulates follicle dormancy(*26, 27*), this localization suggests a potential role in reserve regulation.

Five vascular clusters defined endothelial (Sp-Cl19), smooth muscle (Sp-Cl9, Sp-Cl21), and mural cell populations (Sp-Cl11, Sp-Cl12), supported by GO enrichment (**Supplementary Fig. 2a, Supplementary Table 5**).

Age-dependent differences were evident. Adult cortex was spatially stratified: activated cortical stroma (Sp-Cl6), enriched for cytokine regulation and absent in pediatric tissue, dominated the superficial cortex before transitioning to cortical fibroblasts (Sp-Cl4) at greater depth (**Fig. 3e–f**). As expected, pediatric cortex showed a higher proportion of oocyte-associated signal (Sp-Cl5). Vascular, medullary, and follicle-associated regions were overall more prominent in adult samples, which also contained medulla (**Fig. 3d, Supplementary Fig. 2a,e**).

To further characterize the cortical follicular niche, 137 follicles at comparable developmental stages (primordial and intermediary) were manually annotated, and a 30 µm annular zone extending outward from each oocyte border was defined (**Fig. 3g**). Projection of the 23 Sp-Cls onto these regions revealed marked compositional differences between age groups (**Fig. 3h**). The adult niche was enriched for cortical fibroblasts (Sp-Cl4; ∼2-fold), activated cortical stroma (Sp-Cl6; largely absent in pediatric tissue), and perifollicular stromal myofibroblasts (Sp-Cl17; ∼2.5-fold), whereas the pediatric niche showed higher relative contributions from lymphatic endothelium and macrophages (Sp-Cl13; ∼4.5-fold) and granulosa cells (Sp-Cl14; ∼7.5-fold). The adult perifollicular niche was more homogeneous, dominated by cortical fibroblasts (Sp-Cl4), which comprised ∼50% of the area. In contrast, the pediatric niche was more heterogeneous, containing cortical stroma, interphase, immune, and granulosa populations.

### Transcriptional and structural maturation of the ovarian cortex during puberty

To characterize ovarian maturation, single-cell profiles from chemotherapy-naïve pediatric (n=6) and adult (n=5) ovarian cortex samples were compared. All 14 reference cell types (**Fig. 2b**) were present in both age groups, although their relative abundances and transcriptional profiles differed (**Fig. 4a**).

**Figure 4.**
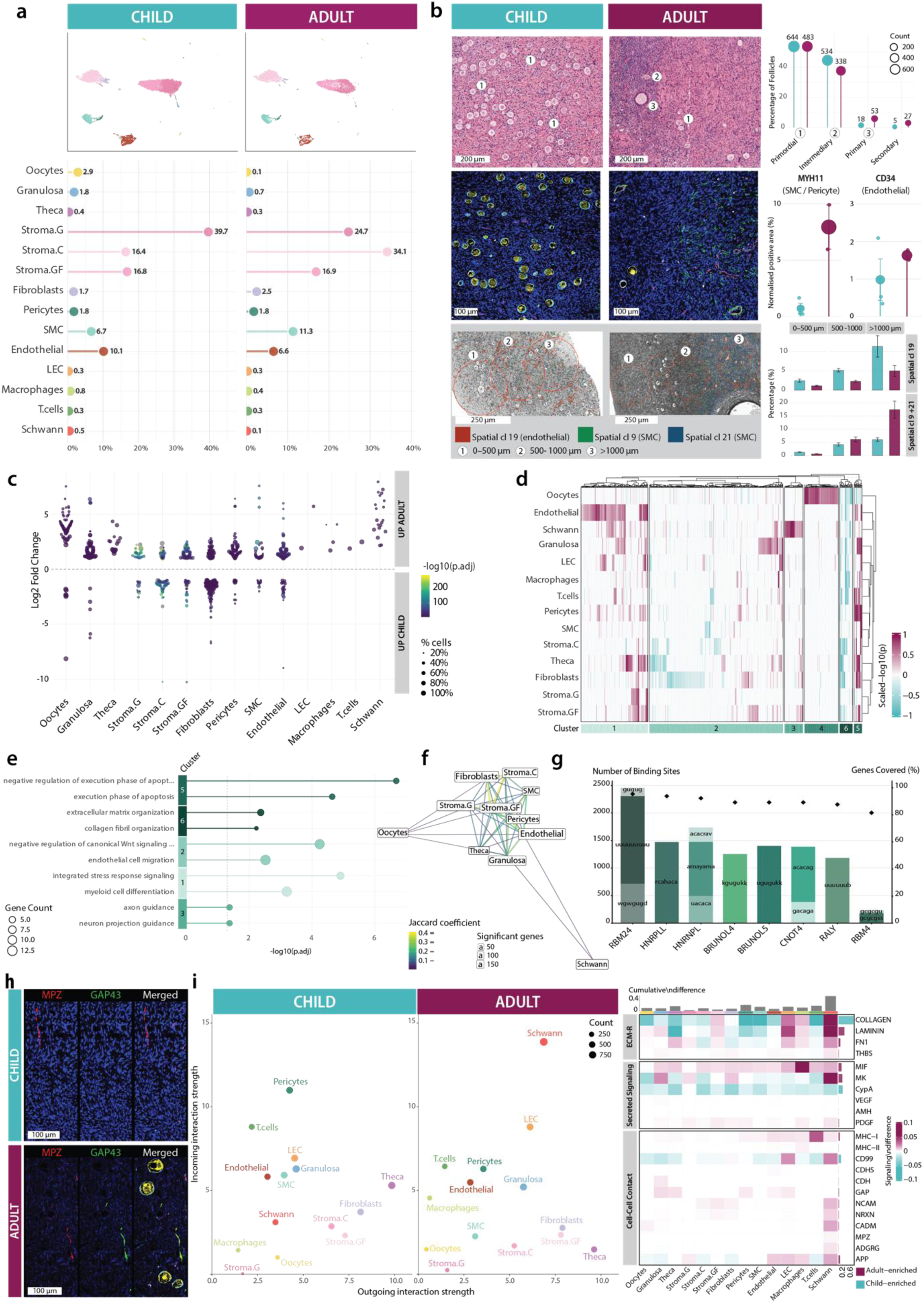
Transcriptional and structural maturation of the ovarian cortex during puberty. **a, UMAP embeddings of single-cell transcriptomic profiles from chemotherapy-naïve child (n = 6) and adult (n = 5) ovarian cortex samples, colored by cell type.** Lollipop plots indicate the relative abundance (%) of the 14 cell types across conditions. **b, Structural characterization of the ovarian cortex across pubertal maturation.** Top, H&E-stained sections of child and adult cortex with follicle stages indicated (① primordial, ② intermediary, ③ primary; secondary follicles not present). Lollipop plot quantifies follicle counts per developmental stage (primordial, intermediary, primary, secondary) in child (n=6) and adult (n=9) samples. Middle, representative images of the mIHC staining panel with MYH11 (smooth muscle cells/pericytes; green) as an additional marker. Plots show normalized MYH11⁺ and CD34⁺ (endothelial cells; magenta) area (%) across ovarian cortex in both chemotherapy-naïve child (n=3) and adult (n=2). Bottom, spatial transcriptomic sections showing endothelial-enriched cluster (Sp-Cl19; red) and SMC–enriched clusters (Sp-Cl9, green; Sp-Cl21, blue) in child and adult tissue. Numbered regions indicate cortical depth zones (① 0–500 µm, ② 500–1000 µm, ③ >1000 µm) with 5 regions of interest per zone per sample analyzed. Bar plots show the percentage area covered by Sp-Cl19 and combined Sp-Cl9 + Sp-Cl21 across depth zones in child (teal) and adult (magenta) **c, Beeswarm plot of DEGs between adult and child samples for each cell type (Wilcoxon rank-sum test; log₂FC > 1, FDR < 0.05).** Each point represents one DEG; the y-axis indicates log₂ fold change (positive values denote upregulation in adult), point size reflects the fraction of expressing cells, and color indicates −log₁₀(adjusted P value). **d, Heatmap of all 574 maturity-associated DEGs across cell types (rows) and genes (columns).** Columns are hierarchically clustered using Ward’s D2 linkage into six transcriptional modules (maturity clusters, Ma-Cl1–6). Rows are ordered hierarchical clustering of cell type transcriptional profiles. Color scale represents signed −log₁₀(P value), with direction indicating upregulation in adult (magenta) or child (teal). **e, GO Biological Process enrichment analysis for Ma-Cls.** Representative terms are shown; dot size indicates gene count and x-axis shows −log₁₀(adjusted P value). Dashed line indicates adjusted P = 0.05. No enrichment was found for Ma-Cl4. **f, Jaccard similarity graph showing the overlap between cell type–specific DEG sets.** Edge width and color represent the Jaccard coefficient; node size reflects the number of DEGs per cell type. Only cell types with at least 10 significant DEGs are shown. **g, RBP binding landscape across adult oocyte-specific genes.** Stacked bars indicate the number of predicted binding sites per motif for selected RBPs. Black diamonds denote the percentage of genes (n = 65) containing at least one predicted binding site. Predictions were derived using RBPmap (Z-score ≥ 4, P ≤ 1 × 10⁻⁴). **h, representative images of the mIHC staining panel with GAP43 (green) as an additional marker in child (top) and adult (bottom) ovarian cortex.** Merged images display the full panel with oocytes (ZP3+GTGSF1+ STAG3) in yellow, granulosa cells (INHA or PRR15) in cyan, blood vessels (CD34) in magenta, immune cells (CD45) in white, and Schwann cells (MPZ) in red. Staining was performed on samples from two children and two adults. **i, Cell–cell communication analysis using CellChat.** Left, scatter plots of outgoing versus incoming interaction strength per cell type in child and adult cortex; point size reflects total interaction count. Right, heatmap of differential signaling pathway activity (adult minus child), grouped by signaling category (ECM–receptor, secreted signaling, cell–cell contact). Magenta indicates adult-enriched signaling; teal indicates child-enriched signaling.

Follicle density declined with age, with follicular cells (granulosa and oocytes) decreasing from 4.7% in children to 0.8% in adults (**Fig. 4a**) (*5*). The granulosa-to-oocyte ratio increased (0.6 to 7), consistent with a higher proportion of growing follicles in adult cortex, as confirmed histologically (**Fig. 4b**). These findings indicate differences in gonadotropin-independent follicle growth dynamics between pediatric and adult cortex.

Rare populations, including immune cells, fibroblasts, and Schwann cells, remained similar in abundance across age groups (**Fig 4a**). In contrast, vascular composition shifted despite similar overall abundance (∼19%), with a decreased endothelial-to-smooth muscle ratio (1.2 to 0.5) indicating vascular maturation. This was supported by mIHC showing increased MYH11-positive smooth muscle coverage and enrichments of spatial smooth muscle-associated clusters (Sp-Cl9, Sp-Cl21) in adults (**Fig. 4b**).

Stromal cell composition also changed, with Stoma-C doubling (16.4% to 34.2%) across puberty, consistent with progressive cortex-medulla segregation (**Fig. 4a, Fig. 3e,f**) (*24, 25*).

Across cell types, puberty was associated with widespread transcriptional remodeling (**Fig. 4c**). Among 574 puberty-associated differentially expressed genes (DEGs), hierarchical clustering identified coordinated gene modules across cell types, with particularly similar programs in stromal, vascular and follicular somatic cell compartments (**Fig 4d-f**).

Two smaller maturity clusters (Ma-Cls) spanning most cell types showed opposite regulation. Upregulated Ma-Cl5 (17 DEGs) was enriched for apoptotic processes driven by *MTRNR2L* pseudogenes encoding humanin-like nuclear isoforms with predicted anti-apoptotic functions. In contrast, downregulated Ma-Cl6 (27 DEGs) represented a stromal program defined by a core matrisome signature, including collagens (*COL1A1, COL1A2, COL3A1*), *DCN* and *FBLN1*, consistent with reduced ECM remodeling in adult cortex (**Fig. 4d,e, Supplementary Tables 6-7, Supplementary Fig. 3a**) (*28*).

Ma-Cl2 (280 DEGs) comprised both up- and downregulated DEGs across most cell types and captured general signaling and migration processes, including negative regulation of canonical WNT signaling pathway and endothelial cell migration. This cluster was driven by stromal genes such as *THBS1*, *LAMA2* and *PODN*, suggesting coordination between ECM and vascular dynamics. Upregulated Ma-Cl1 (140 DEGs) was enriched for integrated stress response signaling and myeloid cell differentiation, with representative DEGs including *CEBPB, JUNB, PPP1R15A*, *MT1A, MT2A*, *MAFB, SOCS1* and *ZBTB16* (**Fig. 4d,e, Supplementary Tables 6-7).**

A distinct oocytes-specific cluster (Ma-Cl4, 71 DEGs) was uniformly upregulated during puberty (**Fig. 4d, Supplementary Table 6**). No enrichment for known GOs, pathways or transcription factor motifs was identified; however, RNA-binding protein (RBP) motifs, including U- and GU-rich elements associated with RBM24, HNRNPL, HNRNPLL, HNRNPA1, and RBM4, were enriched, suggesting coordinated post-transcriptional regulation (**Fig 4g, Supplementary Fig 3b**).

Upregulated Schwann cell-specific Ma-Cl3 (39 DEGs) was enriched for axon guidance and cell migration pathways, including DEGs such as *SEMA3C, GAP43*, and *NCAM1* (**Fig 4d,e, Supplementary Fig. 3a, Supplementary Tables 6-7**). Increased GAP43 expression was confirmed in adult cortex, consistent with pubertal activation of neural guidance programs (**Fig 4h**) (*29*). Across clusters, the selected markers remained similar in prepubertal and pubertal samples, suggesting that the adult phenotype is established only after robust activation of the hypothalamus-pituitary-ovarian axis (**Supplementary Fig. 3a,b**).

Cell–cell communication analysis revealed modest overall changes during puberty but a pronounced increase in Schwann cell activity, particularly in incoming interactions (**Fig. 4i, Supplementary Fig. 3c**). These were dominated by ECM–associated pathways (COLLAGEN, LAMININ, FN1, THBS), together with MK, MIF, and neural signaling (NCAM, NRXN, CADM, MPZ), indicating increased integration of Schwann cells within the cortical niche (**Fig 4i, Supplementary Table 8**). Modest upregulation of cadherin- and gap junction-mediated interactions as well as increased AMH, PDGF, VEGF and MK signaling in granulosa cells suggested enhanced signaling readiness and niche integration (**Fig 4i, Supplementary Table 8**).

Together, these findings demonstrate coordinated structural and molecular maturation of the ovarian cortex during puberty, including vascular remodeling, stromal organization, and broad transcriptional changes across somatic compartments. In parallel, oocytes acquired a distinct post-transcriptionally regulated program, whereas granulosa and Schwann cells adopted a more active and integrated role within the cortical niche.

### First-line chemotherapy triggers extensive transcriptional remodeling of the pediatric ovary

OTC is ideally performed before chemotherapy; however, this is not always feasible. In this cohort, 10 of 16 children had received chemotherapy prior to tissue collection (**Fig. 1c**), enabling assessment of treatment effects across cell types.

Single cell RNA-sequencing data from six chemotherapy-treated children (age 1–13 years) were integrated, yielding 21 transcriptionally distinct clusters (**Fig 5a**). Projection onto the reference atlas (**Fig. 2b**) enabled robust cell type annotation through label transfer, with strong concordance to canonical markers and recovery of most reference populations (**Fig 5a, Supplementary Fig. 4a**). Compared to the naïve pediatric samples (age 7–15 years), chemotherapy was associated with a marked shift in cellular composition, including depletion of vascular populations (pericytes, smooth muscle and endothelial cells) (**Fig. 5b**), supported by mIHC (**Fig. 5c**). Stroma-GF declined from 16.8% to 0.9%, and theca precursors became undetectable. In contrast to depletion of niche populations, oocytes were proportionally increased, likely reflecting the younger patient age (**Fig 1c, Fig. 5b**).

**Figure 5.**
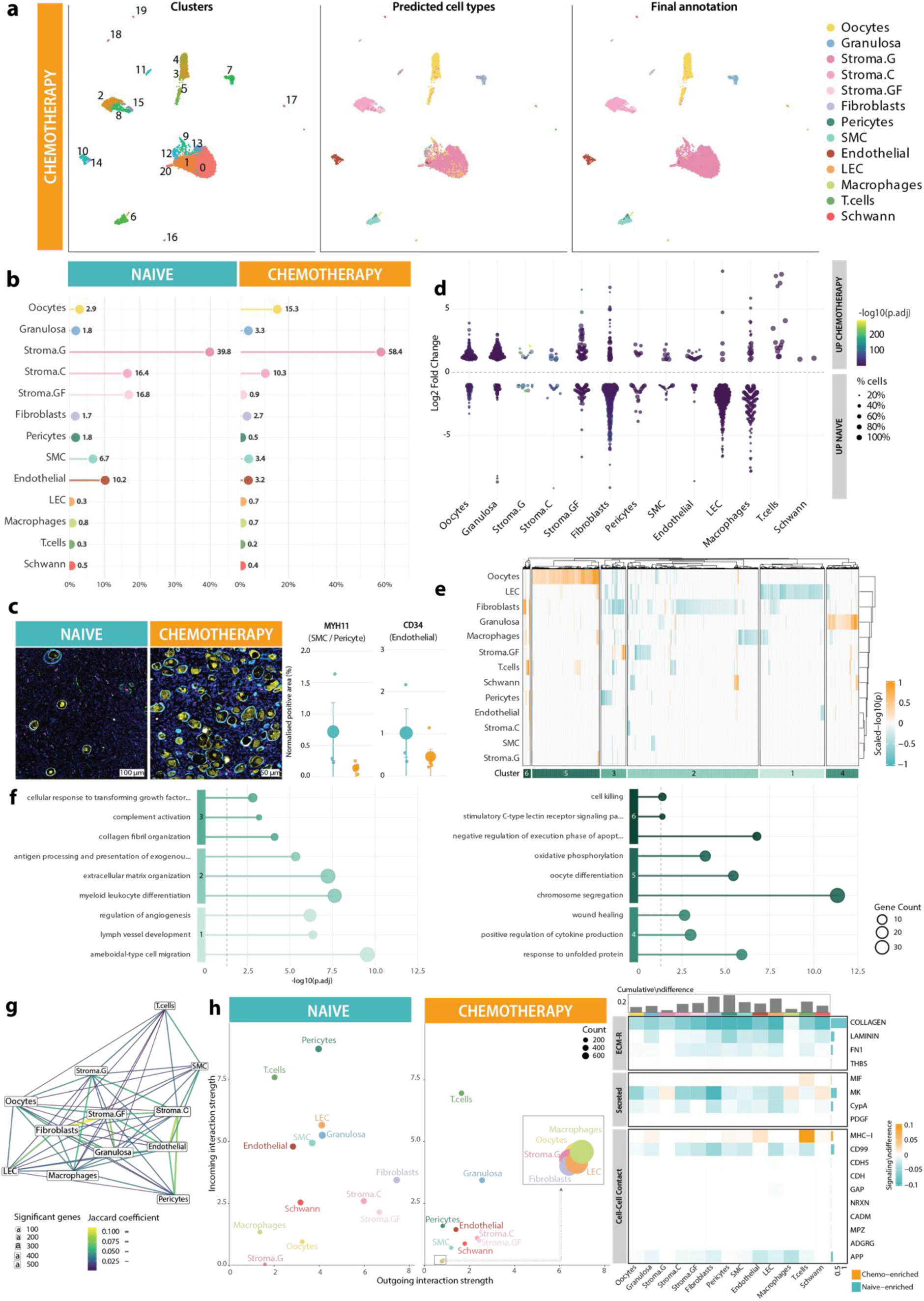
First-line chemotherapy triggers extensive transcriptional remodeling of the pediatric ovary. **a, UMAP projections of single-cell transcriptomes from six chemotherapy-treated pediatric ovarian samples.** Unsupervised clusters (left), predicted cell type identities transferred from the adult/pediatric reference (center), and final cell type annotation (right). Thirteen major ovarian cell populations were recovered. **b, Cell type proportions in chemotherapy-naïve (left) and -treated (right) samples.** Proportions are displayed as percentage of total sequenced cells per condition. **c, Representative images of the mIHC staining panel with MYH11 as an additional marker in naïve (left) and chemotherapy-treated (right) pediatric ovarian cortex.** The full panel shows oocytes (ZP3+GTGSF1+ STAG3) in yellow, granulosa cells (INHA or PRR15) in cyan, blood vessels (CD34) in magenta, immune cells (CD45) in white, and Schwann cells (MPZ) in red. Lollipop plots show normalized MYH11⁺ (smooth muscle cells/pericytes, green) and CD34⁺ (endothelial cells, magenta) area (%) across ovarian cortex in chemotherapy-treated (n=5) and -naïve (n=3) children. **d, Beeswarm plot of DEGs identified per cell type between chemotherapy-treated and -naïve samples.** Each point represents one DEG; point size reflects the maximum cell fraction expressing the gene; color indicates -log₁₀(adjusted p-value). Positive log₂FC values indicate chemotherapy enrichment. **e, Heatmap of all 1,599 unique DEGs across cell types (rows) and genes (columns).** Columns are hierarchically clustered using Ward’s D2 linkage and split into six transcriptional modules (Chemotherapy clusters Ch-Cl 1–6; color bar, bottom). Rows are ordered by hierarchical clustering of cell type transcriptional profiles. Color intensity reflects signed -log₁₀(adjusted p-value) scaled per gene. Orange: chemotherapy-enriched; teal: naïve-enriched. **f, GO Biological Process enrichment lollipop plots for the six DEG clusters.** Ch-Cl 1–3 (left) capture naïve-enriched stromal-immune–vascular programs; Ch-Cl 4–6 (right) capture chemotherapy-associated stress, metabolic and chromatin maintenance programs. Point size indicates gene count; dashed line marks p-adj = 0.05. **g, Jaccard similarity graph depicting the overlap of DEG sets between cell types.** Node size reflects the number of significant DEGs; edge color and width indicate the Jaccard coefficient. Only cell types with ≥ 10 significant DEGs are shown. **h, Cell-cell communication analysis using CellChat.** Left: Scatter plots of inferred outgoing versus incoming interaction strength for chemotherapy-naïve (left) and chemotherapy-treated (right) samples. Dot size reflects the total number of significant ligand–receptor interactions. The insert on the chemotherapy panel shows a magnified view of the cell types with lower interaction strength. Right: differential signaling heatmap showing cumulative and per-pathway signaling differences between conditions (Chemo − Naive), grouped by signaling database (ECM-Receptor, Secreted signaling, Cell–Cell Contact). Orange: chemotherapy-enriched; teal: naive-enriched.

Differential expression analysis identified 1,599 DEGs across cell types, most of which were downregulated following chemotherapy. Schwann cells showed the fewest and fibroblasts the most transcriptional changes (**Fig. 5d**). Hierarchical clustering defined six transcriptional modules segregating into two main axes: downregulated stromal-immune-vascular programs (Chemo clusters Ch-Cls 1-3) and upregulated oocyte-granulosa programs (Ch-Cls 4-5) (**Fig. 5e,f, Supplementary Table 9**).

Ch-Cl1 (316 DEGs) captured a regressed lymphatic endothelial program (*PROX1*, *FLT4*, *KDR, CDH5)*, whereas Ch-Cl3 (122 DEGs) defined a repressed stromal matrisome module (*COL1A1*, *COL1A2, COL3A1, DCN*, *LUM*) primarily affecting fibroblasts and pericytes (**Fig. 5e-f, Supplementary Tables 9-10**). The large predominantly downregulated Ch-Cl2 (638 DEGs) integrated ECM organization with immune-associated processes, supported by matrix-remodeling and antigen presentation genes (*MMP14, ADAMTS5, HLA-DQB1*).

In contrast, upregulated Ch-Cls 4–6 reflected stress responses, energy metabolism changes, and chromosomal processes (**Fig 5e-f, Supplementary Tables 9-10**). Ch-Cl4 (158 DEGs), associated with granulosa cells, was enriched for stress response pathways including unfolded protein response and cytokine signaling (*ATF3, DDIT3, CHOP, XBP1, IL6ST,* and *SERPINE1)*. Ch-Cl5 (329 DEGs), strongly enriched in oocytes, was defined by chromosome segregation and oxidative phosphorylation, driven by cell-cycle regulators (*AURKA, CCNB1, CCNB2, MAD2L1, UBE2C*), oocyte factors *(FIGLA, GDF9, ZP3*), and mitochondrial genes (*NDUFA6, COX5A*), indicating activation of proliferative and metabolic programs. The small, upregulated Ch-Cl6 (36 DEGs) was enriched for anti-apoptotic and cytotoxic pathways including *MTRNR2L* pseudogenes, *GZMH*, and *KLRC4* (**Fig 5e-f, Supplementary Tables 9-10**).

Classical chemotherapy-induced cytotoxicity, apoptosis and inflammatory responses (*BBC3, BAX, TP53, IL6, CXCL8, TNF*) were not detected, consistent with resolution of acute effects at the time of sampling (**Supplementary Fig. 4b**). Despite low pairwise Jaccard coefficients for DEG overlaps between cell types, transcriptional connections were present across most compartments (**Fig. 5g**), indicating a widespread but context-dependent chemotherapy response rather than a specific shared gene signature.

Cell–cell communication analysis revealed a ∼70% collapse in intercellular signaling following chemotherapy (**Fig 5h, Supplementary Fig. 4c, Supplementary Table 11**). ECM-associated pathways (COLLAGEN, LAMININ, FN1, THBS) and secreted signaling through MK were markedly attenuated across cells. In granulosa cells, cadherin- and gap-junction mediated interactions were lost alongside AMH and PDGF signaling (**Supplementary Table 11**). Only a limited number of non-ECM pathways, notably MHC-I signaling by T-cells, showed relative enrichment within an otherwise suppressed signaling landscape in the chemotherapy-treated child ovaries (**Fig 5h, Supplementary Table 11**).

Collectively, chemotherapy depleted vascular and follicular niche populations, reduced ECM production and associated signaling, and induced stress-associated programs in follicles.

The post-chemotherapy pediatric ovary adopts a damage-adapted, signaling-suppressed state.

### Chemotherapy targets the pubertal transcriptional program in the pediatric ovary

Given the prominent role of matrisome-related pathways in both ovarian maturation and the response to chemotherapy, the corresponding transcriptional programs were compared. Overall, 56% of puberty-associated DEGs were also affected by chemotherapy (**Fig. 6a**). Shared upregulated genes (n=131, 31% of the puberty signature) were enriched for stress response signaling, vascular development, and cell cycle regulation, whereas shared downregulated genes (n=132, 57%) were strongly associated with ECM organization, cell–substrate adhesion, focal adhesion, and cell junctions (**Fig 6a, Supplementary Table 12**). To further compare transcriptional response across cell types, pairwise correlation analysis of log₂ fold-change signatures was performed across 27 cell type–condition combinations. Oocytes, granulosa cells, stroma-C, and endothelial cells exhibited strongly coordinated responses between puberty and chemotherapy (Pearson r > 0.5) (**Supplementary Fig. 5**). Network and hierarchical clustering analyses identified these compartments as having the most closely related transcriptional responses (**Fig 6b**). Functional enrichment analysis of concordantly regulated genes per cell type showed enrichment of growth factor signaling, stress responses, cellular proliferation and differentiation pathways among upregulated genes, whereas downregulated genes were associated with ECM production, collagen fibril organization, and cytoskeletal dynamics (**Fig. 6c, Supplementary Table 13**). Across all cell types, concordantly responses were associated with increased stress and vascular remodeling signatures together with reduce ECM organization and cell adhesion pathways (**Fig. 6d, Supplementary Table 13**).

**Fig. 6.**
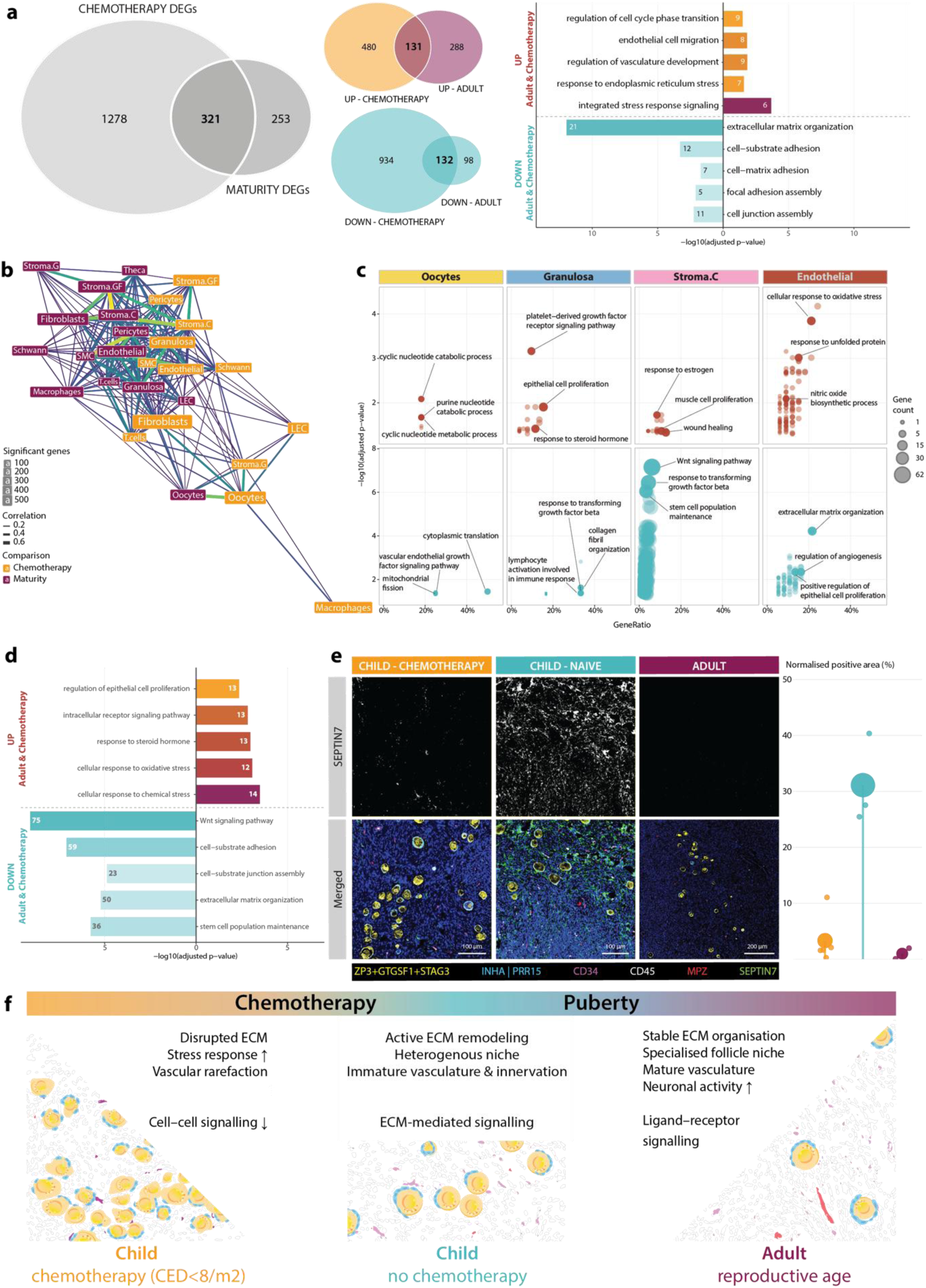
Chemotherapy targets the pubertal transcriptional program in the pediatric ovary. **a, Overlap of DEGs associated with puberty (Adult vs Child) and chemotherapy exposure (Chemotherapy vs Naïve).** Venn diagrams show shared and condition-specific overall (left) and downregulated/upregulated (right) genes. Right, GO Biological Process enrichment of shared DEGs. Bar length indicates −log10 adjusted P value; numbers denote gene counts. **b, Cross-condition transcriptional similarity across cell types**. Pairwise Pearson correlations of log₂ fold-change signatures (permissive threshold) are shown as a network (edges: r > 0.1; edge width/color proportional to correlation strength). **c, GO Biological Process enrichment of concordantly regulated genes in highly correlated cell types (r > 0.5).** Upregulated (red) and downregulated (teal) programs are shown separately, highlighting shared pathways across conditions. Dot size represents gene count. **d, Global GO enrichment analysis of pan-cell-type overlap gene sets, showing shared biological processes across all cell types.** **e, SEPTIN7 expression across conditions (chemotherapy-treated and -naïve child, and adult).** Left, representative images of the mIHC panel with SEPTIN7 (green) as an additional marker. Merged images display the full panel with oocytes (ZP3+GTGSF1+ STAG3) shown in yellow, granulosa cells (INHA or PRR15) in cyan, blood vessels (CD34) in magenta, immune cells (CD45) in white, and Schwann cells (MPZ) in red. Right, lollipop plot shows pixel-classified SEPTIN7-positive area per sample (chemotherapy-naïve child, n = 3; chemotherapy-treated child, n = 5; adult, n = 2). **f, Schematic model summarizing condition-specific structural and signaling states in the ovarian cortex.** Chemotherapy at cumulative doses CED < 8/m^2^ induces ECM disruption, stress responses, vascular rarefaction, and collapse in cell-cell signaling; the chemotherapy-naïve pediatric ovary is characterized by active ECM remodeling, immature vasculature, and a heterogeneous niche; and adult tissue exhibits stable ECM architecture, specialized follicular niches, mature blood vessels and innervations, with active ligand–receptor–mediated signaling.

To identify candidate biomarkers associated with these changes, septin family genes emerged as candidates of interest given their established roles in tissue morphogenesis and mechanical integrity through linking the plasma membrane to the cytoskeleton (**Supplementary Table 13**) (*30*). Immunohistochemical analysis revealed marked differences in SEPTIN7 expression across puberty and in response to chemotherapy (**Fig. 6e**). Chemotherapy-naïve pediatric ovarian cortex exhibited robust and widespread SEPTIN7 staining, whereas signal intensity was reduced following chemotherapy and largely absent in adult tissue (**Fig 6e**). These findings support a role for SEPTIN7 in maintaining mechanical integrity of the ovarian cortical niche, which appears sensitive to both chemotherapy and pubertal maturation.

Collectively, the results support a model in which chemotherapy response co-opts key elements of puberty-associated transcriptional remodeling while driving a maladaptive state marked by ECM disorganization, impaired cell–cell communication, and destabilization of septin cytoskeletal architecture (**Fig. 6f**).

## DISCUSSION

Gonadotoxic therapies affect ∼ 14,000 children annually in Europe, and improved survival has created a growing population at risk of infertility and other long-term sequelae (*31*). In females, high-dose alkylating agents and radiotherapy compromise ovarian function (*1, 31*), yet current understanding of ovarian toxicity relies largely on clinical endpoints and histology, providing limited insights into cell type-specific targets. Direct investigation of the human ovary, particularly in children, remains rare. Here, clinically relevant pediatric ovarian tissue was systematically profiled to resolve cellular architecture, pubertal maturation, and responses to chemotherapy.

Adult samples showed vascular maturation in the cortex, reflected by a thicker MYH11+ smooth muscle layer (*32*), and activation of the cortical neural compartment, evidenced by increased GAP43 expression and enhanced nerve-associated adhesion signaling in Schwann cells. As vascular and neural structures extend from the medulla into the cortex and support follicle growth (*29, 33*), these findings suggest progressive establishment of an integrated cortical niche during puberty. Spatial analysis further demonstrated a more structured cortical organization in adults, with increased representation of specialized myofibroblasts surrounding cortical follicles, whereas the pediatric niche remained more heterogeneous. In parallel, granulosa cells of the adult ovarian reserve showed increased paracrine signaling, including MK and AMH pathways, together with strengthened cadherin-and gap junction-mediated cell-cell interactions. Together, these findings suggest that ovarian cortical maturation involves development of a more functionally specialized microenvironment that may support the greater growth capacity of gonadotropin-independent follicles observed in adult ovarian tissue (*12, 13*).

Parallel to niche maturation, the transcriptional landscape of ovarian reserve oocytes also changed across puberty. Adult oocytes upregulated genes involved in chromatin regulation and genome stability (*SMARCB1, LEO1, HENMT1*), ribosome biogenesis (*BRIX1, LTV1, BYSL*), and cell cycle control (*CKS1B, KIF18A, HMMR*). This maturation germline program was enriched for motifs of RBPs involved in post-transcriptional regulation (*34, 35*). RBPs have also been identified as key regulators of oocyte competence in goat (*36*) and linked to male infertility in humans (*37*). As human oocytes rely heavily on post-transcriptional regulation during maturation (*38*), these findings suggest that RBP-mediated control may already be established in primordial follicles during puberty, potentially contributing to the enhanced genome regulation and translational capacity of the adult ovarian reserve. This is consistent with known chromatin reorganization during oocyte maturation (*39*) and the reduced developmental competence of prepubertal oocytes, characterized by impaired meiotic fidelity (*40*), and weaker translational programs (*41*). The germline program in the three pubertal patients remained similar to the prepubertal state, suggesting that the acquisition of full germline readiness occurs after activation of the hypothalamic–pituitary–ovarian axis.

Alkylating agents target both proliferating and quiescent cells, explaining the vulnerability of dormant follicles (*42*). In our samples, collected weeks after first-line chemotherapy, no chemotherapy-related increase in canonical apoptosis, cell death, or inflammation markers was detected, likely reflecting resolution of acute responses (*43*). Yet, chemotherapy induced markedly greater perturbation than puberty, with depletion of stroma-GF, theca precursor, and vascular populations, together with widespread transcriptional remodeling. Vasculature is an established target of alkylating agents (*44*), and disruption of the somatic niche likely further compromises follicle growth potential. Cell–cell communication was reduced by ∼ 70%, including loss of MHC class II–related signaling. In granulosa cells, lost AMH signaling (*45*) along with gap junction and cadherin interactions suggest impaired follicle integrity and signaling capacity.

One of the most unexpected findings was the substantial overlap between puberty- and chemotherapy-associated transcriptional changes, particularly in ECM-related pathways. Across somatic cell types, puberty was associated with reduced expression of collagens, decorin and lumican, consistent with previous proteomic studies (*28, 46*), potentially affecting tissue mechanobiology and availability of growth factors, which could link to the increased readiness for TGF-β, VEGF and PDGF signaling in adult samples. Notably, 57% of genes downregulated during puberty were also downregulated following chemotherapy exposure. The overlapping transcriptional program was enriched for ECM organization, cell junctions, and cell-matrix adhesion, particularly coordinated in follicular, cortical stromal and endothelial compartments. These signatures suggest loss of ECM regulation and associated signaling (TGFβ, VEGF). ECM remodeling has been implicated in major ovarian transitions including puberty and menopause, as well as in disease states such as PMOS (*11, 47–49*). Interestingly, puberty and menopause timing are also influenced by shared genetic variants (*50*). These findings suggest that ovarian maturation, chemotherapy response, and reproductive aging may share similar, biologically malleable features, for example related to tissue remodeling. However, despite partial transcriptional similarities, chemotherapy does not recapitulate physiological ovarian maturation, as no functional niche or follicle growth competence is established.

Biomarkers of chemotherapy-induced damage could improve quality assessment of ovarian tissue collected for fertility preservation. While classical acute damage markers such as PUMA, p53, and BAX(*51–53*) were not detected at the time of tissue collection, SEPTIN7 expression differed markedly across developmental and treatment states, possibly serving as a longer-lasting tissue-state biomarker. Given the role of septins in cytoskeletal organization and mechanical integrity (*30, 54*), we hypothesize that septins participate in niche changes during both puberty and chemotherapy exposure.

Our study has limitations, including small sample size and heterogeneity in diagnoses, ages and treatment regimens. Its strength lies in the analysis of freshly collected pediatric ovarian tissue within a clinical fertility preservation setting, reflecting real-world material collected for potential future transplantation and fertility restoration.

Our findings have direct clinical implications. Puberty drives coordinated maturation of both the somatic compartment and germline, establishing a spatially organized niche that supports follicle growth competence. In contrast, chemotherapy disrupts cortical cell populations and signaling, including loss of niche interactions. Consequently, immature ovarian tissue, particularly after chemotherapy exposure, may have limited capacity to fully recapitulate adult maturation and function following transplantation. Fertility preservation should therefore, whenever possible, be performed before treatment initiation or with minimal chemotherapy exposure, while strategies to support tissue maturation *in vitro* warrant further exploration.

Whether the child ovary can recover from chemotherapy-induced changes remains to be determined in longitudinal studies. The capacity of re-transplanted tissue to support fertility is determined by both the ability of prepubertal tissue to complete normal maturation and the extent of chemotherapy-induced damage (*22*). As such, OTC in pre-pubertal children should still currently be considered as investigational.

## METHODS

### Sveafertil study – collection of research samples and metadata

The Sveafertil study was set up following the Nordic Society of Paediatric Haematology and Oncology and Swedish national recommendations for fertility preservation in children (*23*). Patients aged 1–17 years scheduled to receive gonadotoxic treatment associated with a high (postmenarche) or very high (premenarche) risk of infertility were eligible. In premenarcheal patients, very high risk was defined as direct ovarian irradiation (>10 Gy) or planned allogeneic or autologous hematopoietic cell transplantation. In postmenarcheal patients, high risk was defined by the same criteria, with the addition of a cumulative CED exceeding 6–8 g/m². Exclusion criteria included increased risk of infection, bleeding, pain, or other surgical complications. Written age-adapted information was provided to families and to children ≥7 years of age. Informed consent was obtained from legal guardians and assent from the child when applicable. The Sveafertil study was approved by the Swedish Ethical Review Authority (2019–03802). Additional prepubertal samples collected within a fertility preservation research project at Helsinki University Hospital, Finland, using identical inclusion and exclusion criteria, were included under separate ethical approval (340/13/03/03/2015). All research samples were pseudonymized and stored in registered biobanks.

Ovarian tissue collection was coordinated with a clinically indicated surgical procedure requiring general anesthesia. Approximately one-third of one ovary was excised; two-thirds of that biopsy was cryopreserved for potential future clinical use and one-third allocated for research within the Sveafertil study. At the time of surgery, one vial of venous blood was collected for hormonal assessment. Research samples were transported to Karolinska Institutet in DPBS supplemented with calcium and magnesium (Thermo Fischer Scientific) using dedicated temperature-controlled containers (2–8 °C) with continuous tracking. Transport time ranged from <1 h to up to 20 h. Upon arrival, tissue was immediately processed for single-cell transcriptomic analysis or fixed in 4% methanol-free formaldehyde for spatial transcriptomics and multiplex immunostaining, or in Bouin’s solution for histological assessment. In cases of larger biopsies, additional tissue fragments were cryopreserved using established controlled-rate freezing protocols (*55*). Clinical meta-data included age, diagnosis, and treatment protocol. Cumulative CED presented alkylating agent exposure(*56*). Anthracycline exposure was calculated as cumulative doxorubicin isotoxic equivalent dose (DIE) using established conversion factors: 1.0 for doxorubicin, 0.5 for daunorubicin, 5.0 for idarubicin, and 10.0 for mitoxantrone (*57*).

Samples used in this work were collected during years 2020 – 2024.

### Serum hormone analysis and pubertal staging

Available serum samples were analyzed for luteinizing hormone (LH), estradiol (E2), follicle-stimulating hormone (FSH), and anti-Müllerian hormone (AMH) at the Clinical Chemistry Laboratory, Karolinska University Hospital. Pubertal status in patients aged >7 years was defined by Tanner stage and serum thresholds for hypothalamic–pituitary–gonadal axis activation (LH > 0.3 IU/L; E2 > 80 pmol/L) (*58*).

### Adult sample collection

Adult ovarian control samples were obtained from individuals undergoing elective abdominal surgeries (gender-affirming surgery or caesarean section). Participants received written study information and provided informed consent prior to donation of ovarian tissue and peripheral blood. In gender-affirming surgery, tissue that would otherwise have been discarded was donated in its entirety. In elective caesarean section cases, a superficial cortical biopsy (≤5 mm diameter, ≤2 mm thickness) was collected. Metadata included age, body mass index, smoking status, and relevant diagnoses or treatments with potential impact on ovarian function. All samples and data were pseudonymised and stored in a registered biobank. Tissue was retrieved directly from the operating theatre and transported to the laboratory within 15 min in DPBS supplemented with calcium and magnesium (Thermo Fischer Scientific) for immediate processing into single-cell libraries or formalin- or Bouin-fixed paraffin-embedded blocks, as described above for pediatric samples. The study was approved by the Swedish Ethical Review Authority (2015/798-31, with amendments).

### Histological assessment

All samples underwent histological quality control. Bouin-fixed, paraffin-embedded tissue was serially sectioned (4 µm), H&E-stained, and digitized using a Pannoramic Scanner (3DHistech, Budapest, Hungary). Every tenth section was evaluated using Pannoramic Viewer software (v1.15.4, 3DHistech) or QuPath (v0.7.0) for identification of follicles within the cortex. For follicle stage identification, one section per patient was analyzed. Primordial and primary (unilaminar) follicles were identified as oocytes surrounded by a single layer of flattened or cuboidal granulosa cells, while intermediary follicles were surrounded by a mix of these cells. Secondary follicles had at least two layers of cuboidal granulosa cells.

### Single-cell RNA sequencing

Fresh ovarian cortical tissue was mechanically and enzymatically dissociated into single-cell suspensions immediately upon arrival in the laboratory. Libraries were prepared using the Chromium Single Cell 3’ platform (v2 chemistry, 10x Genomics). Dissociation protocols were optimized during the study to accommodate differences in tissue texture between pediatric and adult samples: first three samples were processed according to our previously published protocol(*59*), and subsequent samples were dissociated using a GentleMACS device with a Tumor Dissociation Kit (Miltenyi). One sample was processed using a modified dissociation protocol consisting of Collagenase IV (Thermo Scientific), Liberase TM (Sigma-Aldrich), and DNase (Qiaqen) in DMEM/F12 supplemented with FBS, followed by incubation at 37°C for 40 min in a MACS Tube Rotator. Cell suspensions were filtered (35-40 μm cell strainer, Miltenyi Biotec), counted using an automated cell counter (Moxi, Orflo Technologies), and assessed for viability by trypan blue exclusion prior to loading onto the Chromium controller for GEM generation. Cell input varied between 2,000 to 10,000 cells per sample, with viability ranging 79-97%. Libraries were sequenced on an Illumina NextSeq 500 platform.

### Single-cell RNA-seq data processing and analysis

#### Data preprocessing

Raw sequencing data were processed using Cell Ranger v6 (10x Genomics) to generate feature–barcode count matrices. Individual samples were imported into Seurat (v.5.2.1) and subjected to quality control filtering, retaining cells with 200–5,000 detected genes, <30,000 unique molecular identifier (UMI) counts, and <20% mitochondrial transcript content.

#### Integration

Following filtering, a total of 17 ovarian cortex samples were included, comprising 6 chemotherapy-naïve pediatric samples, 6 chemotherapy-exposed pediatric samples, and 5 adult samples. To establish a reference dataset of the unperturbed ovarian cellular landscape, chemotherapy-naïve pediatric and adult samples (n = 11) were initially integrated.

Data were normalized using Seurat’s default log-normalization method, and the 2,000 most highly variable genes were identified using variance-stabilizing transformation (VST). Scaled expression values were used for principal component analysis (PCA). To correct for sample-level batch effects, Harmony was applied using sample identity as the grouping variable. Cross-condition integration was subsequently performed using reciprocal PCA (RPCA) as implemented in Seurat v5, using Harmony-corrected embeddings as input. Cells were clustered using a shared nearest-neighbor (SNN) graph-based approach with a resolution of 0.9, based on the first 30 integrated dimensions. Low-dimensional visualization was performed using Uniform Manifold Approximation and Projection (UMAP).

#### Cluster annotation

Cluster annotation was performed using a multi-evidence approach. Differentially expressed genes per cluster were identified using the Wilcoxon rank-sum test as implemented in Seurat (log∼2∼ fold-change threshold = 0.25, minimum cell fraction = 0.20, positive markers only). Cell type identity was assigned based on canonical marker gene expression and confirmed by GO overrepresentation analysis using clusterProfiler (v.4.10.1) (Biological Process, Molecular Function, and Cellular Component ontologies; BH-adjusted p < 0.01). Automated predictions (**Supplementary Table 14**) were additionally generated using GPT-4 via the GPTCelltype framework, queried with ovarian cortex-specific tissue context. Reference-based annotation was independently performed using SingleR (v.2.4.1) against three curated references: the Human Primary Cell Atlas, the Blueprint/ENCODE compendium, and the Database of Immune Cell Expression (DICE). Cluster marker genes were further cross-referenced against the CellMarker 2.0 database, restricted to human normal ovarian cell types. Final cell type identities were assigned by integrating these orthogonal sources with manual curation and stroma subtype identities were further supported by spatial expression patterns. Spatial deconvolution was performed at an 8 µm resolution using Semla (v1.1.6), integrating Visium HD data with matched single-cell RNA sequencing reference profiles. This approach enabled estimation of cell type composition within spatial bins, leveraging the complementary strengths of high-resolution spatial data and single-cell transcriptomic annotations.

Five clusters were excluded prior to final annotation (clusters 6, 14, 16, 19, and 23) based on patient-specific enrichment, low abundance, or expression profiles consistent with non-ovarian reproductive tract contamination. The remaining clusters were annotated into 14 cell type categories, with identities validated by GO Biological Process overrepresentation analysis per cell type using clusterProfiler; three representative terms per cell type were selected for visualization.

#### Maturity analysis (Adult vs Child)

DEGs between adult and chemotherapy-naive pediatric ovarian samples were identified per cell type using the Wilcoxon rank-sum test as implemented in Seurat (log₂ fold-change threshold = 1, minimum cell fraction = 0.20, two-sided). Genes with a Benjamini–Hochberg adjusted p-value < 0.05 were considered significant. DEGs were visualized as a beeswarm plot displaying log₂ fold-change per cell type, with point size proportional to the maximum cell fraction and color encoding −log₁₀(adjusted p-value).

To identify shared transcriptional programs across cell types, significant DEGs were aggregated into a gene × cell type matrix of signed –log10(P values), linearly rescaled and capped at the 95th percentile per cell type and hierarchically clustered using Ward’s D2 linkage into six gene clusters. Cell types with no result for a given gene were assigned a value of 0. Pairwise overlap between cell type DEG sets was quantified using the Jaccard similarity coefficient, restricted to cell types with at least 10 significant DEGs. For each gene cluster, the top enriched GO Biological Process terms (Benjamini–Hochberg adjusted p-value < 0.05) were computed using clusterProfiler (v4.10.1); a representative subset of terms was manually selected per cluster and visualized as a lollipop plot.

Vascularization differences were studied using spatial transcriptomic results from child and adult ovaries, analyzed using QuPath (v0.5.1). High-resolution H&E-stained images of spatially analyzed samples were loaded together with spatial cluster information provided as GeoJSON files. Regions of interest (ROIs) were defined as circular annotations with a diameter of 500 µm. Because the adult samples covered a larger area, the maximum depth of the child samples was used to standardize the analysis and set the corresponding limit for adult samples. Three ROIs were defined: 0-500 μm (1), 500-1000 μm (2), and >1000 μm (3). Endothelial cells were identified using Sp-Cl19, while smooth muscle cells were identified using Sp-Cl9 and Sp-Cl21. The percentage of cells within each ROI was calculated using a QuPath Groovy script.

Genes differentially expressed in adult oocytes were analyzed for enrichment of RBP motifs. Genomic regions corresponding to 65 protein-coding genes were retrieved using the biomaRt package (v2.64.0), subdivided into overlapping 10-kb windows, and scanned for known human and mouse RBP motifs using RBPmap (v1.3) with the default background model. The 25 most significantly enriched motifs (z-score > 4, P < 1 × 10⁻⁴) were selected for independent validation. Consensus motifs were converted to MEME-compatible formats and scanned against the same genomic regions using FIMO (MEME Suite). Zero-order and first-order Markov background models were generated from the nucleotide composition of the 65 genes and applied during FIMO analysis. Motifs with significant occurrences (P < 1 × 10⁻⁴) across genomic segments and at the gene level were aggregated to confirm motif enrichment across the gene set.

Ligand–receptor interactions were inferred using CellChat (v2.1.2) with the combined signaling database (secreted signaling, ECM-receptor, and cell–cell contact). Separate CellChat objects were constructed for adult and chemotherapy-naive pediatric samples and compared to identify condition-enriched interactions. Differential interaction counts and strengths were visualized as heatmaps, and overall signaling activity per pathway was summarized as the mean of outgoing and incoming communication probabilities. A curated subset of pathways grouped by signaling category was additionally examined.

#### Chemotherapy effect analysis

Six chemotherapy-treated ovarian cortex samples were pre-processed independently. Following the same normalization and dimensionality reduction pipeline applied to the reference dataset (i.e. log-normalization, identification of the 2,000 most variable genes by VST, scaling, and PCA) sample-level batch effects were corrected using Harmony with sample identity as the grouping variable, followed by RPCA integration to further harmonize the query dataset internally. Clustering was performed on a shared nearest-neighbor graph at resolution 0.9 using the first 30 integrated dimensions, and a UMAP embedding was computed for quality control purposes. Sample and treatment group contributions to each cluster were assessed using stacked bar plots

The pre-processed chemotherapy-treated dataset was subsequently projected onto the Adult/Child reference dataset. Transfer anchors between the reference and query were identified using reciprocal PCA, and cell type probabilities were propagated to each query cell via label transfer. The query was additionally embedded in the reference UMAP space to enable direct visual comparison with the reference annotation.

Cluster marker genes were identified using the Wilcoxon rank-sum test (log2 fold-change threshold = 1, minimum cell fraction = 0.20, positive markers only; BH-adjusted p < 0.05). Marker expression was cross-referenced against canonical ovarian cell type signatures using dot plot visualization.

Prediction confidence was assessed per cell type as the median label transfer score. Cluster purity, defined as the fraction of cells assigned to the dominant predicted cell type, was computed from the cluster × predicted cell type contingency table (**Supplementary Table 15**). GO enrichment analysis (Biological Process ontology) was performed per predicted cell type using clusterProfiler (BH-adjusted p < 0.05), restricted to cell types with at least 10 marker genes.

Final cell type labels were assigned by manual curation integrating predicted identities, marker expression, and cluster purity metrics. Two clusters required sub-resolution: cluster 6, containing a mixture of smooth muscle and perivascular cells, and cluster 16, containing a mixture of T cells and macrophages, were resolved by computing lineage-specific module scores, assigning each cell to the identity with the highest score. Although theca precursor cells were predicted by label transfer, they were recovered in low numbers and did not form a discrete cluster; they were therefore not assigned as a separate cell type in the final annotation.

Downstream analyses analogous to those performed for the maturity comparison were applied to the chemotherapy effect dataset. Briefly, significant DEGs (Benjamini–Hochberg adjusted p-value < 0.05) were aggregated into a signed −log₁₀(p-value) matrix, hierarchically clustered into six gene clusters (Ward’s D2), and subjected to GO Biological Process enrichment analysis using clusterProfiler (v4.x), with a curated subset of terms visualized as lollipop plots. Pairwise DEG overlap between cell types was quantified using the Jaccard similarity coefficient (restricted to cell types with ≥ 10 significant DEGs). Ligand–receptor interactions were inferred using CellChat with the combined signaling database, and differential interaction counts and strengths between chemotherapy-treated and naive samples were visualized as heatmaps and pathway-level summaries.

#### Overlap between puberty- and chemotherapy-associated transcriptional programs

To identify shared transcriptional changes between puberty and chemotherapy exposure, DEG sets computed independently for each condition (see above) were intersected per direction: upregulated overlap genes were defined as genes significantly upregulated in both the Adult vs Child and Chemotherapy vs Naïve comparisons; downregulated overlap genes as those significantly downregulated in both. Intersections were computed at the global level and independently per shared cell type. GO Biological Process enrichment of the global upregulated and downregulated overlap gene sets was performed using clusterProfiler (v4.10.1) with Benjamini–Hochberg correction (adjusted p < 0.05, q < 0.2); a curated subset of enriched terms was selected for visualization.

#### Cross-condition transcriptional similarity and identification of convergent cell-type programs

To assess similarity between puberty- and chemotherapy-associated transcriptional changes across ovarian cell types, we computed pairwise Pearson correlations between log₂ fold-change vectors. Differential expression was re-estimated using a permissive threshold (|log₂FC| > 0.1, Benjamini–Hochberg-adjusted P < 0.05, minimum cell fraction = 0.10; Wilcoxon rank-sum test) to capture broader transcriptional programs. Genes with zero fold-change in either condition were excluded.

Correlations across the 27 cell type–condition combinations (14 maturity, 13 chemotherapy) were compiled into a symmetric matrix and visualized as a network (edges at r > 0.1; edge weight and color scaled by correlation strength) and a hierarchically clustered heatmap (Euclidean distance, Ward’s D2). Network layout was generated using the Fruchterman–Reingold algorithm (igraph v2.1.1; ggraph v2.2.1).

Cell types with Pearson r > 0.5 between maturity- and chemotherapy-associated profiles were defined as exhibiting convergent transcriptional responses. For these cell types, overlap gene sets were defined as genes showing concordant regulation (up or down) in both conditions at the permissive threshold (adjusted P < 0.05). GO Biological Process enrichment was performed separately for up- and downregulated genes using clusterProfiler (v4.10.1; org.Hs.eg.db v3.18.0), with Benjamini–Hochberg correction (adjusted P < 0.05, q < 0.2; minimum gene set size = 3).

To obtain a global view of shared processes, pan-cell-type overlap gene sets were generated by taking the union of concordantly regulated genes across all cell types and analyzed as above.

### Visium HD spatial transcriptomics

#### Tissue preparation and imaging

Formalin-fixed paraffin-embedded (FFPE) ovarian biopsy samples from three children (aged 3–8 years) and three adults (aged 22–24 years) were sectioned at 4 µm on Superfrost Plus slides. Sections were deparaffinized and rehydrated through xylene, absolute ethanol, 96% (v/v) ethanol, 70% (v/v) ethanol, and tap water, followed by haematoxylin staining for 3 min, a tap water rinse, and eosin staining for 1 min. Sections were then placed in nuclease-free water before mounting with 85% (v/v) glycerol for imaging. High-resolution H&E images were captured at 20× magnification using the Metafer Slide Scanning platform (microscope stand, AxioImager.Z2 with ScopeLED illumination, Zeiss; camera, CoolCube 4m, MetaSystems; objective, Plan-Apochromat 20×/0.80 M27, a = 0.55 mm, Zeiss; software, Metafer5 version 3.14.192). Raw images were stitched with VSlide software (v1.1.128; MetaSystems).

#### Library preparation

Following imaging, slides were de-coverslipped, destained, and de-crosslinked as described in the Visium HD FFPE Tissue Preparation Handbook (CG000684 Rev B, 10x Genomics). Probe hybridization, probe ligation, Visium HD slide preparation, probe release, extension, pre-amplification, SPRIselect cleanup, and library construction were performed as outlined for the Visium HD slide with 6.5 × 6.5 mm capture area in the Visium HD Spatial Gene Expression Reagent Kits User Guide (CG000685 Rev A, 10x Genomics).

#### Sequencing

Visium HD libraries contain standard Illumina paired-end constructs beginning with P5 and ending with P7. Spatial barcodes are encoded at the start of TruSeq Read 1 (Read 1T), while i7 and i5 sample index sequences are incorporated as the index reads. Read 1T was used to sequence the 43 bp spatial barcode and UMI, and Small RNA Read 2 (Read 2S) was used to sequence the ligated probe insert. Libraries were sequenced on the Illumina NextSeq 2000 platform using a P3 200-cycle flow cell. Minimum sequencing depth was calculated as the product of the estimated tissue coverage percentage and 275 million read pairs per fully covered capture area, as recommended by 10x Genomics for FFPE tissues. This minimum threshold was met and exceeded for all samples.

#### Visium HD data processing

Raw FASTQ files were processed using Space Ranger (v3.0.0; 10x Genomics) with the GRCh38 human reference transcriptome (refdata-gex-GRCh38-2020-A). H&E images were manually aligned to the Visium HD capture area within Space Ranger using the Loupe Browser manual alignment wizard. Gene expression matrices were generated at 2 µm and 8 µm resolutions.

#### Cell segmentation

Oocytes were identified using a hybrid transcriptomic and image-based strategy. Due to their large size relative to surrounding cell types, standard segmentation approaches often fragment oocytes into multiple objects corresponding to subcellular structures. To address this, we first performed coarse spatial aggregation by binning the data into 8 µm grids and conducted clustering followed by differential expression analysis to identify oocyte-enriched regions. These regions were projected onto the corresponding H&E images and used as spatial priors for segmentation with Segment Anything Model (v1.0; SAM). SAM was applied using the identified oocyte-enriched bins as prompts to delineate oocyte boundaries in the histological images.

In instances where adjacent oocytes were merged into a single segmentation object, a custom post-processing procedure was applied. This procedure leveraged the morphological properties of oocytes—specifically their approximately circular geometry and high area-to-perimeter ratio—to identify and separate merged objects into individual oocytes using shape-based splitting heuristics.

Segmentation of all remaining cells was performed using StarDist (v0.9.1), a deep learning–based method optimized for star-convex object detection in microscopy images. To avoid double counting, segmentation was restricted to regions outside the previously defined oocyte boundaries. This ensured that oocyte regions were exclusively represented by the SAM-derived masks, while all other cell types were captured by StarDist.

#### Transcript-to-cell assignment and quality control

Spatial transcriptomic bins at 2 µm resolution were assigned to segmented cells based on geometric consistency. Specifically, each bin was assigned to a cell if all four vertices of the bin polygon had their nearest boundary points associated with the same segmented cell. This conservative assignment strategy minimized ambiguity at cell boundaries and reduced cross-contamination of transcripts between adjacent cells.

Cells with fewer than 25 UMIs were excluded from downstream analyses. Highly variable genes were identified using scanpy (v1.11.4), and the top 5,000 were selected for modelling.

#### Dimension reduction and clustering

Dimensionality reduction was performed using scVI (v1.3.3), a deep generative model that accounts for technical variability and sparsity in single-cell and spatial transcriptomics data. A latent space of 10 dimensions was learned and used for downstream analyses. Clustering was conducted using the Leiden algorithm implemented in standard single-cell analysis frameworks, with a resolution parameter of 1.1 to identify spatially coherent transcriptional clusters.

#### Differential expression analysis

Differential expression analysis between spatial clusters was performed using PyDESeq2 (v0.5.2), which models UMI counts as a negative binomial distribution and accounts for dispersion estimation and normalization. For each spatial cluster, cells assigned to that cluster were compared against all remaining cells (one-vs-rest). Wald tests applied to log-fold change estimates and *P-*values were adjusted using the Benjamini-Hochberg procedure; genes with adjusted *P <* 0.05 and |log2 fold-change| > 1 were considered differentially expressed.

#### Cortex segmentation and cortical analysis

In pediatric samples, the entire tissue section was considered as cortex due to the small size of these surface biopsies. For adult samples, which were derived from ovarian cross-sections containing both cortex and medulla, the cortex was operationally defined as the superficial 1-mm-thick layer extending inwards from the cortical surface towards the medulla.

To determine the tissue boundary, we first identified the largest contiguous area within the slide that was not covered by tissue, corresponding to background space in the histological image. The interface between this background region and the tissue was taken as the outer tissue boundary. All spatial positions within the tissue whose Euclidean distance to this boundary was less than 1 mm were classified as belonging to the cortical region.

This geometric definition provided a consistent and reproducible approximation of the cortical compartment across samples, independent of local variations in tissue morphology.

To characterize the depth-dependent spatial organization of the cortex, the 1 mm cortical region was subdivided into 50 concentric strips of 20 µm width, oriented parallel to the outer tissue boundary, with strip 1 corresponding to the outermost cortical surface. For each strip, the total area occupied by cells assigned to each spatial cluster was calculated and expressed as spatial cell-cluster area per strip (µm²).

#### Perifollicular niche analysis

To investigate the cellular microenvironment surrounding oocytes, a subset of cortical follicles (n = 77 pediatric, n = 60 adult; total n= 137) was selected by manual inspection of the H&E images. This selection was performed to retain only high-confidence primordial and intermediary follicles with clearly defined unilaminar morphology and to exclude cases where oocytes were in close proximity to one another, which could confound spatial neighborhood analyses.

For each selected oocyte, a perifollicular neighborhood was defined as an annular region extending radially 30 µm outward from the oocyte boundary, excluding the oocyte itself. All cells whose centroids fell within the specified radial distance from the oocyte boundary were included in that neighborhood. Using the cell-segmentation masks obtained as described above, the contribution of each spatial cluster was quantified as its share of the total cell area within the perifollicular zone.

### Multiplex immunostaining

The mIHC panel was built to target five ovarian structures and cell populations: oocytes (cocktail of three antibodies: GTSF1, HPA038877, 1:800; STAG3, HPA049106, 1:4000; and ZP3, HPA054061, 1:2000), granulosa cells (INHA, CAB079772, 1:100; or PRR15, HPA040996, 1:500), immune cells (PTPRC/CD45, HPA000440, 1:200), endothelial cells (CD34, CAB000018, 1:250), and Schwann cells (MPZ, HPA068925, 1:2000), plus one open slot for any protein or structure of interest. Detailed information on all antibodies is provided in **Supplementary Table 16**. Six tyramide-linked OPAL fluorophores (OPAL 480, 520, 570, 620, 690, and TSA-DIG/780; Akoya Biosciences, Marlborough, MA) were used and each OPAL fluorophore was paired with its respective antibody based on expected co-expression and relative abundance to minimize crosstalk and optimize signal balance, i.e., low-expressing markers were paired with high-intensity fluorophores. Antibodies for the five fixed markers in the panel were selected and matched with a suitable OPAL dye based on standard HPA workflow as previously described (*60*). The staining cycle order and OPAL dyes were combined as follows: granulosa cells – 690, immune cells – 620, open slot – 520, Schwann cells – 570, oocytes – 480, and endothelial cells – TSA-DIG/780. FFPE ovary tissue blocks were sectioned at 4 µm, dried overnight at RT, and baked at 50 °C for 12-24 h. Before staining, slides were processed following previous protocols (*60*). In brief, slides were deparaffinized in xylene and rehydrated in graded alcohols (99.9%, 96%, and 80%) with blocking of endogenous peroxidase activity using 0.3% hydrogen peroxide. Slides were washed under running deionized water and then subjected to heat-induced epitope retrieval (HIER) using a Decloaking chamber (BioCare, CA, USA) at 125 °C for 4 min in pH 6 Dako Target Retrieval Solution Citrate (DAKO, Glostrup, Denmark). The same pH 6 buffer was used for every HIER step except the last one where pH 9 (Thermo Fisher Scientific, USA) was used instead. Slides were cooled passively to 90 °C and placed under running deionized water for 1 min and stored in TBS buffer supplemented with 0.2% (v/v) Tween 20 (Immunologic, Arnhem, The Netherlands). mIHC was performed by iterative staining-stripping cycles with the Opal 6-plex Detection Kit for Whole Slide Imaging (Akoya Biosciences, Marlborough, MA, USA) as follows: slides were first blocked with Peroxidase Block (Cell Marque, 925B-04), then primary antibodies were incubated for 30 min at RT, then incubated with secondary reagents by using HiDef Detection HRP Polymer System (Cell Marque, CA, USA) for 10 min, and finally OPAL fluorophores for 10 min. Between each incubation step, slides were washed in TBS supplemented with 0.2% (v/v) Tween 20 (Immunologic, Arnhem, The Netherlands). After each staining round, antibody complexes were denatured by HIER at 90 °C for 20 min, and a new round of antibody staining was performed. OPAL fluorophores were reconstituted in DMSO and diluted 1:500 in 1X Plus Automation Amplification Diluent (Akoya Biosciences). Following staining, slides were treated with Ready Probes Tissue Autofluorescence Quenching kit (#R37630, Thermo Fischer Scientific, Eugene, Oregon, USA), counterstained with DAPI (1:1000, Invitrogen, D1306, Thermo Fisher Scientific) and mounted with ProLong Glass Antifade mounting medium (ThermoFisher Scientific, MA, USA). Whole-slide multispectral images were acquired on the Akoya PhenoImager at with 40x objective. Digital qptiff images were acquired for downstream quantitative and spatial analysis.

Multiplex immunohistochemistry images were analyzed using QuPath (v0.6.0). For cross-sectional adult tissue specimens, a ROI corresponding to the outermost 1 mm of the ovarian cortex was delineated programmatically using a custom Groovy script. For all other tissue types (chemotherapy-naïve pediatric samples, chemotherapy-treated pediatric samples, and adult cortical biopsies), the entire tissue area was considered as the ROI and was manually annotated.

Marker positivity within each ROI was quantified using per-sample pixel classifiers applied at moderate resolution (2 µm/pixel for MYH11 and CD34) or low resolution (4 µm/pixel for SEPTIN7). Intensity thresholds were manually adjusted for each tissue section to account for inter-sample variability in staining and image acquisition. Positive pixel area was expressed as the percentage of total tissue area within the ROI above the marker-specific threshold.

## Statistical analysis

No statistical method was used to predetermine sample size. The experiments were not randomized. The Investigators were not blinded to allocation during experiments and outcome assessment.

## Data availability

Processed single-cell data, along with raw and processed spatial sequencing data generated in this study, are available in the ArrayExpress repository under accession numbers E-MTAB-16786 and E-MTAB-16938, respectively. All mIHC images generated in this study will be made publicly available upon publication through Figshare (https://doi.org/10.6084/m9.figshare32111734.).

## Code availability

All code used for data processing and analysis in this study will be made publicly available upon publication at https://github.com/DamdimopoulouLab/Pubertal-maturation-and-chemotherapy-associated-disruption-of-the-paediatric-ovary, including scripts and documentation necessary to reproduce the analyses.

## Acknowledgements

We thank the Sveafertil network and Reproductive Medicine Karolinska for patient recruitment and sample handling. We acknowledge the Morphological Phenotype Analysis facility (FENO) for histological slide preparation and scanning, and the Human Protein Atlas (HPA) team in Uppsala for their contributions. Imaging was performed at the Live Cell Imaging Core Facility/Nikon Center of Excellence at Karolinska Institutet, supported by the KI Infrastructure Council. Bioinformatics support was provided by the BEA (Bioinformatics and Expression Analysis) core facility at NEO, supported by the Karolinska Institutet Board of Research and the Research Committee at Karolinska University Hospital. We also acknowledge the National Genomics Infrastructure (NGI), Sweden, for sequencing support, and Biobank and Study Support at Karolinska University Hospital for their professional service. We thank Drs Per Frisk, Cecilia Petersen, Ravindra Naraine, Ludvig Bergenstråhle, Patrick Truong, Fei Luo, and Marco Vicari for valuable input and assistance.

This work was supported by grants from the Swedish Research Council (2020–02132, 2024–02647), the Swedish Childhood Cancer Fund (2020-02132, 2024-02647, KP2023-0030), Karolinska Institutet (SFO, KID, Consolidator Grant), State research funding of Helsinki University Hospital, the Foundation for Paediatric Research, and the Finnish Cancer Society.

## CRediT author contributions

**Conceptualization:** PD, KJ, LM, RM

**Methodology:** PD, AD, LM, RM, MWM, JH, KK, CLi, HT

**Software:** HT

**Validation:** CLi, AD

**Formal analysis:** LM, HT, MWM, JH, IR, MA, NB, AD

**Investigation:** PD, RM, LM, JH, FB, MWM, EP, KK, AJ, BK, RS, MO, Cli

**Resources:** PD, RM, KPe, HV, KPa, MS, PB, CLi, JM, TT, JL, GG, Cla

**Data curation:** PD, KJ, RM, LM, JH, FB, IR, MWM, HT

**Writing – original draft:** PD, KJ, LM, MWM, JH

**Writing – review & editing:** PD, KJ, RM, LM, JH, IR, MWM, NB, AJ, HV, KPa, MS, PB, CLi, HT, KPe, AS, JM, TT, JL, GG, CLa

**Visualization:** LM, AD, HT, MWM, JH, IR, FB, NB, FH

**Supervision:** PD, KJ, RM, CLi

**Project administration:** PD, RM, JH, KPa, CLi

**Funding acquisition:** PD, TT, KJ

## Abbreviations

CED: cyclophosphamide equivalent dose
Ch-Cl: chemotherapy cluster
DEG: differentially expressed gene
ECM: extracellular matrix
GO: gene ontology
H&E: haematoxylin-eosin
Ma-Cl: maturity cluster
mIHC: multiplex immunohistochemistry
OTC: ovarian tissue cryopreservation
RBP: RNA binding protein
ROI: region of interest
Sp-Cl: spatial cluster
UMI: unique molecular identifier.

## Notes

### Competing Interest Statement

The authors have declared no competing interest.

