## Supplementary Figures for "Pubertal maturation and chemotherapy-associated disruption of the pediatric ovary revealed by multimodal single-cell profiling"

#### Supplementary Figure 1

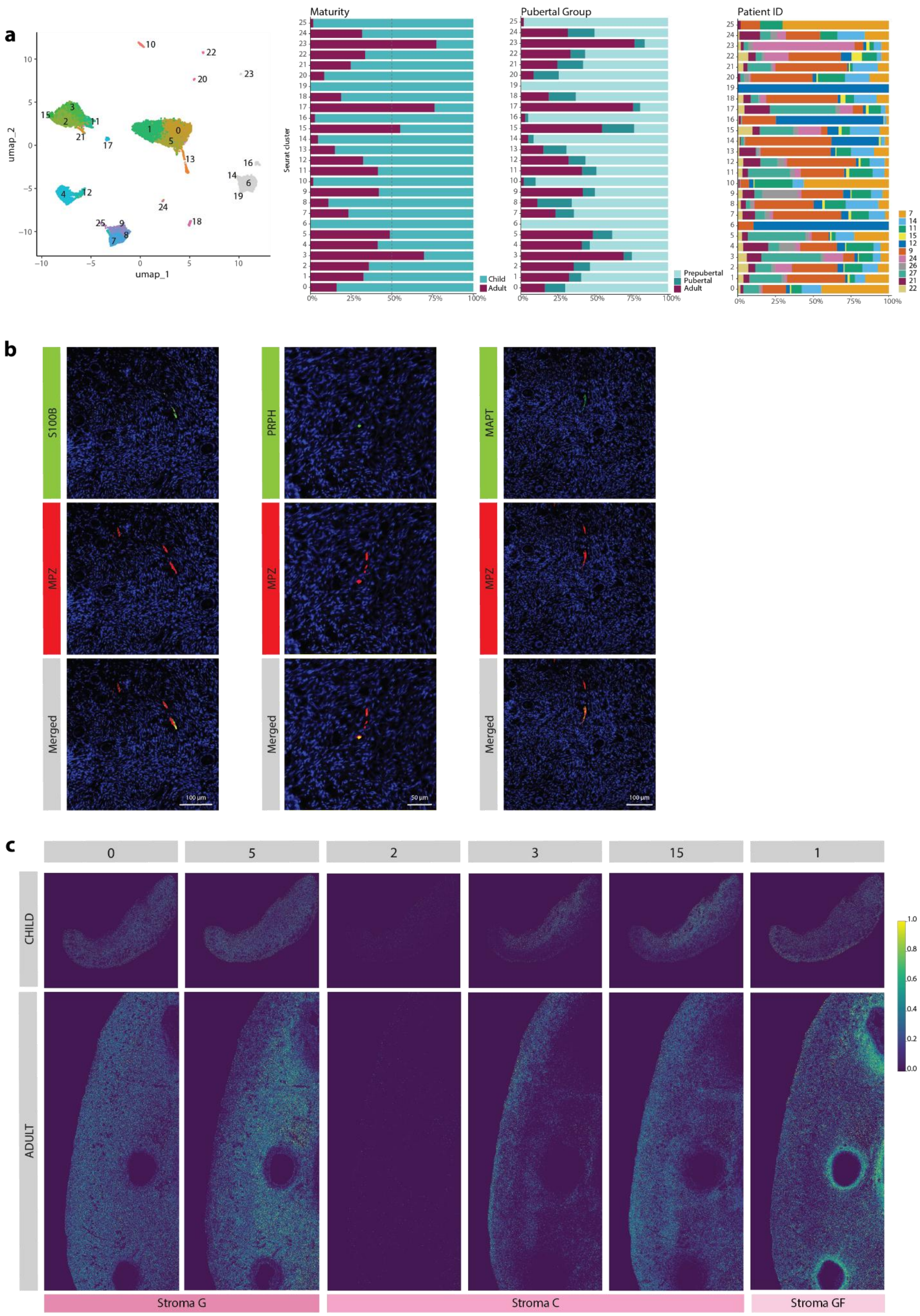

**Supplementary Figure 1. | Integration, validation, and spatial characterization of ovarian cortex cell populations.**
a, UMAP visualization of integrated single-cell transcriptomes from treatment-naïve pediatric (6–16 years, n=6) and adult (25–32 years, n=5) ovarian cortex samples, showing initial clustering into 26 populations. Clusters excluded from downstream analysis due to batch effects (clusters 6, 14, 16 and 19) or reproductive tract contamination (cluster 23) are shown in grey. Bar plots indicate the proportional contribution of each cluster by sample maturity, pubertal group, and individual patient, demonstrating robust integration across donors with minimal batch effects aside from the excluded clusters.
b, Multiplex immunohistochemistry (mIHC) validation of Schwann cells in ovarian cortex. Representative images show co-localization of the Schwann cell marker MPZ (red) with S100B, PRPH and MAPT (green), with merged channels confirming spatial overlap. Nuclei are counterstained with DAPI (blue). Staining was performed on one representative adult sample.
c, Spatial deconvolution mapping of stromal subpopulations. Heatmaps from one pediatric and one adult sample show the spatial distribution of stromal subtypes derived from original Seurat clusters: stroma-G (sc-clusters 0 and 5), stroma-C (sc-clusters 2, 3 and 15), and stroma-GF (sc-cluster 1). Signal intensity reflects the relative expression of cluster-defining genes, highlighting spatial compartmentalization in stromal organization.

#### Supplementary Figure 2

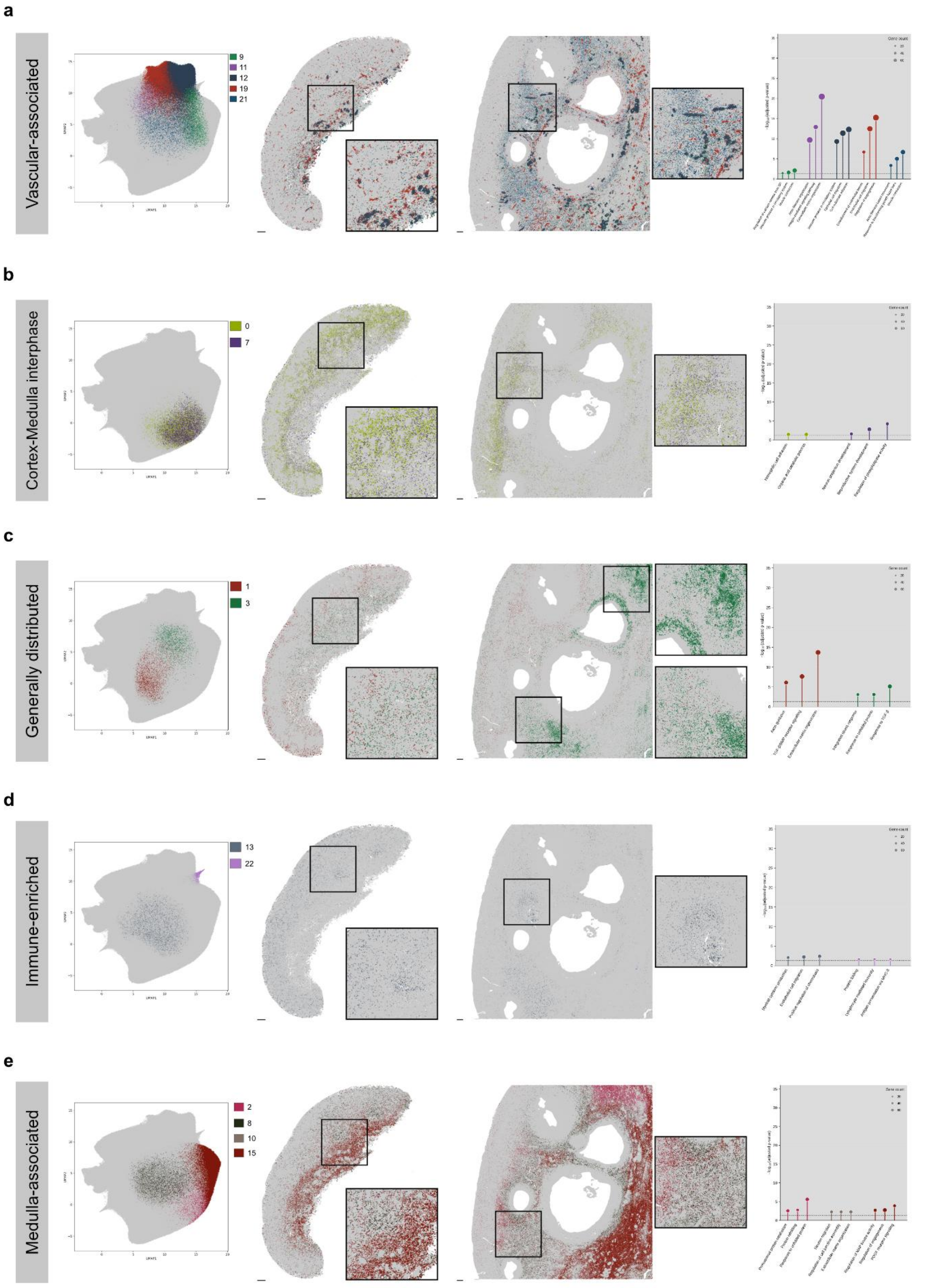


**Supplementary Figure 2. | Spatial localization and gene ontology enrichment of remaining compartments.**

a, Vascular-associated compartment. Left, UMAP highlighting vascular-associated clusters with remaining clusters greyed out. Middle, spatial localization on representative child (Patient 3, left) and adult (Patient 17, right) tissue sections, with magnified insets. Right, lollipop plot showing GO enrichment; the y-axis shows $-\log_{10}$(adjusted p-value) and dot size represents gene count. Dashed line indicates the significance threshold (adjusted P = 0.05).
b, Cortex–medulla interphase compartment. Left, UMAP highlighting interphase clusters with remaining clusters greyed out. Middle, spatial localization on representative child (Patient 3, left) and adult (Patient 17, right) tissue sections, with magnified insets. Right, lollipop plot showing GO enrichment, displayed as in (a).
c, Generally distributed compartment. Left, UMAP highlighting generally distributed clusters with remaining clusters greyed out. Middle, spatial localization on representative child (Patient 3, left) and adult (Patient 17, right) tissue sections, with magnified insets. Right, lollipop plot showing GO enrichment, displayed as in (a).
d, Immune-enriched compartment. Left, UMAP highlighting immune-enriched clusters with remaining clusters greyed out. Middle, spatial localization on representative child (Patient 3, left) and adult (Patient 17, right) tissue sections, with magnified insets. Right, lollipop plot showing GO enrichment, displayed as in (a).
e, Medulla-associated compartment. Left, UMAP highlighting medulla-associated clusters with remaining clusters greyed out. Middle, spatial localization on representative child (Patient 3, left) and adult (Patient 17, right) tissue sections, with magnified insets. Right, lollipop plot showing GO enrichment, displayed as in (a).
Scale bars: 200 μm throughout.

### Supplementary Figure 3

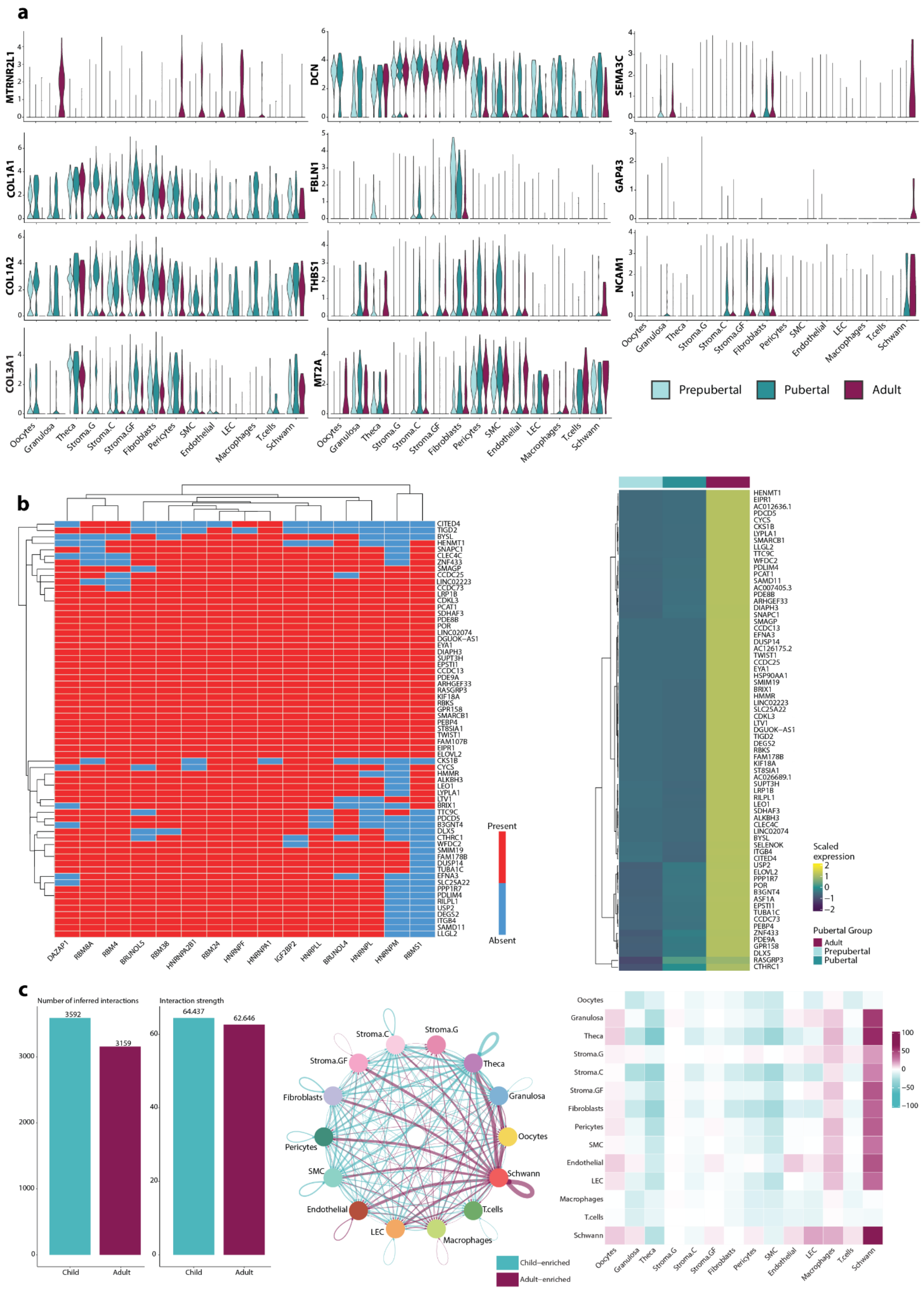

**Supplementary Figure 3. | Transcriptional dynamics of the maturing human ovary across pubertal development and cell–cell communication remodeling.**
a, Violin plots showing the expression of selected candidate genes across cell types, split by pubertal group (Prepubertal, Pubertal, Adult). Genes are grouped by maturity transcriptional cluster: Ma-Cl 5 (MTRNR2L family; apoptosis-related, upregulated in Adult), Ma-Cl 6 (*COL1A1, COL1A2, COL3A1, DCN,* and *FBLN1*; pediatric-enriched matrisome, downregulated in Adult), Ma-Cl 2 (*THBS1*; signaling and stromal-associated, upregulated in Adult), Ma-Cl 1 (*MT2A*; stress response signaling, upregulated in Adult), and Ma-Cl 3 (*SEMAC3*, *GAP43*, and *NCAM1*; Schwann cells specific, upregulated in Adult).
b, Left, binary heatmap showing the presence or absence of RNA-binding protein (RBP) motifs (FIMO analysis) across oocyte DEGs. Right, heatmap of scaled average expression of oocyte Ma-Cl 4 genes (n=71) across pubertal groups (Prepubertal, Pubertal, Adult), with hierarchical clustering (Ward's D2). Color scale represents row-scaled expression (z-score).
c, Left, bar plots showing the total number of inferred cell–cell interactions (left) and cumulative interaction strength (right) in Adult and Child conditions. Centre, circle plot depicting differential interaction strength between Adult and Child; magenta edges indicate Adult-enriched interactions, teal edges indicate Child-enriched interactions. Right, heatmap of differential interaction counts per sender–receiver cell type pair (Adult minus Child).

### Supplementary Figure 4

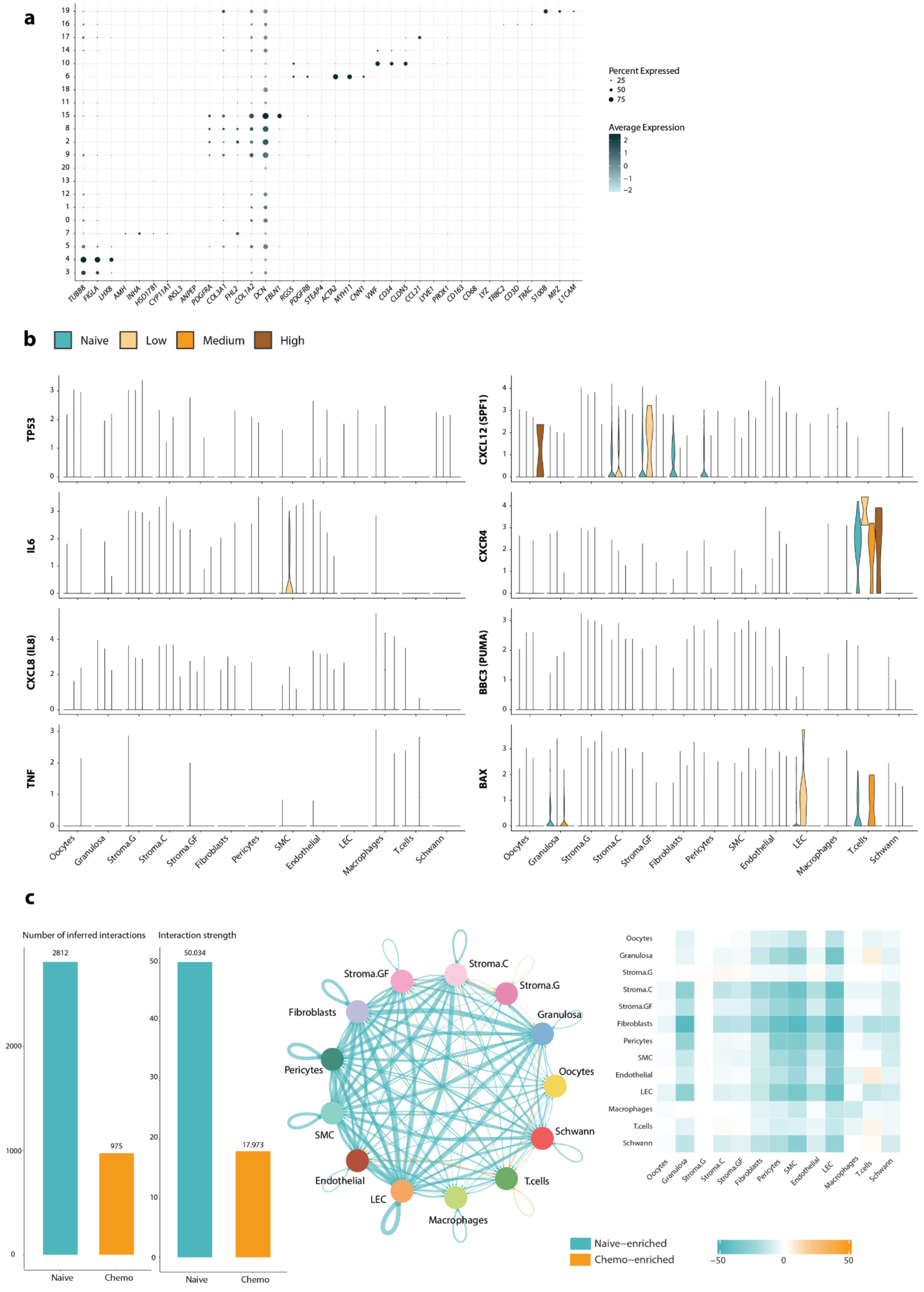


**Supplementary Fig. 4 | Cell-type annotation, chemotherapy response signatures and altered cell–cell communication after chemotherapy.**

a, Dot plot showing expression of canonical marker genes across non-annotated cell clusters in chemotherapy-exposed pediatric ovarian samples. Dot size indicates the percentage of cells expressing each gene and color indicates average expression.
b, Violin plots showing expression for classical chemotherapy-associated cytotoxicity, apoptosis and inflammatory response signatures across ovarian cell types, stratified by chemotherapy exposure category: treatment-naïve, low, medium and high.
c, Cell–cell communication analysis comparing treatment-naïve and chemotherapy-exposed pediatric ovarian samples. Left, bar plots showing the total number of inferred interactions and cumulative interaction strength in treatment-naïve and chemotherapy-exposed conditions. Centre, circle plot depicting differential interaction strength between conditions; teal edges indicate naïve-enriched interactions and orange edges indicate chemotherapy-enriched interactions. Right, heatmap showing differential interaction counts per sender–receiver cell-type pair.

#### Supplementary Figure 5

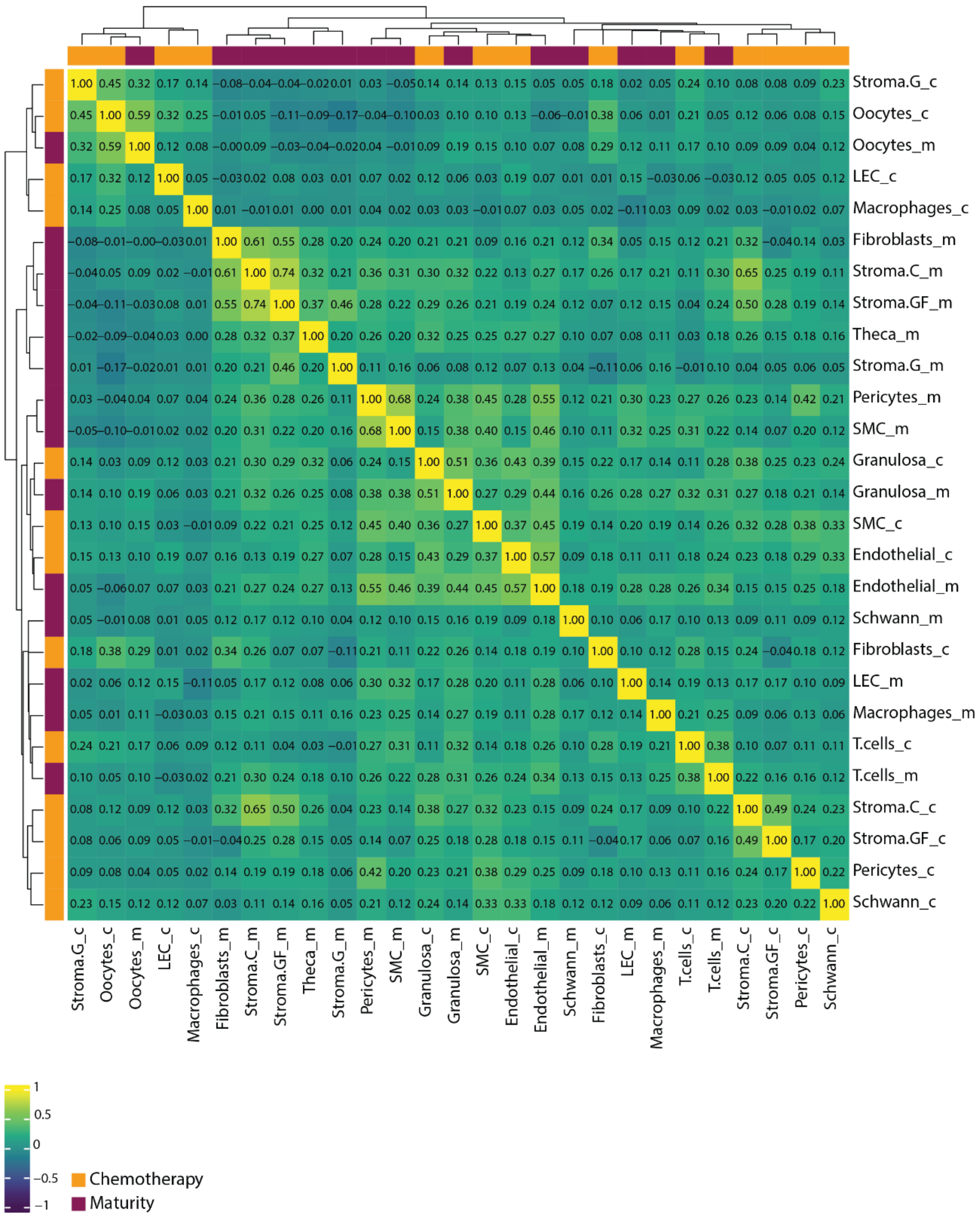

**Supplementary Figure 5. | Pairwise Pearson correlation matrix of cell-type transcriptional responses to puberty and chemotherapy.**
Symmetric heatmap of pairwise Pearson correlations between log2 fold-change vectors from the Maturity (Adult vs Child, _m; magenta) and Chemotherapy (Chemo vs Naive, _c; orange) comparisons across all shared ovarian cell types. Fold-change vectors were computed at a lenient threshold ($|\log2FC| > 0.1$, Benjamini–Hochberg-adjusted $p < 0.05$) to capture the full transcriptional signature of each condition. Genes with zero fold-change in either condition were excluded. Rows and columns are ordered by hierarchical clustering (Euclidean distance, Ward D2 linkage).
